# Redox regulation of autophagy in Arabidopsis: differential effects of reactive oxygen species

**DOI:** 10.1101/2023.08.11.552961

**Authors:** Germán Robert, Alejandro Enet, Laura Saavedra, Ramiro Lascano

## Abstract

Autophagy plays a key role in the responses to different stress condition in plants. Reactive oxygen species (ROS) are common modulators of stress responses, having both toxic and signaling functions. In this context, the relationship between ROS and autophagy regulation remains unclear, and in some aspects, contradictory. In this study, we employed pharmacological and genetic approaches to investigate the effects of different ROS on the cytoplastic redox state and autophagic flux in *Arabidopsis thaliana*. Ours results demonstrated that oxidative treatments with H_2_O_2_ and MV, which drastically increased the oxidized state of the cytoplasm, reduced the autophagic flux. Conversely, singlet oxygen, which did not have significant effects on the cytoplasmic redox state, increased the autophagic flux. Additionally, our findings indicated that after H_2_O_2_ and high light treatments and during the recovery period, the cytoplasm returned to its reduced state, while autophagy was markedly induced. In summary, our study unveils the differential effects of ROS on the autophagic flux, establishing a correlation with the redox state of the cytoplasm. Moreover, it emphasizes the dynamic nature of autophagy in response to oxidative stress and the subsequent recovery period.

**HIGHLIGHTS:**

- This research shows the differential effects of reactive oxygen species on autophagic modulation, highlighting their impact on the cytoplasmic redox state. The relationship between ROS and autophagy regulation remains unclear, and in some aspect’s contradictory. Here, we present a comprehensive investigation characterizing the effects of different ROS, such as hydrogen peroxide and singlet oxygen, on the modulation of autophagy in Arabidopsis. In brief, our findings reveal differential impacts on cytoplasmic redox states and autophagic flux, providing insight into the dynamic nature of autophagy, especially in stress and post-stress conditions.
- To the best of our knowledge, our work is the first to evaluate autophagic flux both during and after oxidative stress in plants. Our results indicate that this differentiation is crucial when analyzing the effects of oxidative stress on autophagy.

## INTRODUCTION

Plants as sessile organisms are constantly exposed to diverse stress conditions, triggering mechanisms that integrate defense and acclimation responses (Fichman and Mittler, 2020; Peláez-vico *et al*., 2022). A common effect of stress is the increased production of reactive oxygen species (ROS), such as superoxide (O_2_ ^-^), hydrogen peroxide (H O), singlet oxygen (^1^O), and the hydroxyl radical (-OH), at different subcellular compartments, such as chloroplasts, mitochondria, peroxisomes, cytosol and the apoplast (Apel and Hirt, 2004; Foyer and Noctor, 2009, 2011*a*). Depending on their site, intensity, and duration of production, ROS can modify biomolecules, influence redox signaling, and drive processes like growth, senescence, and cell death. Their levels are tightly regulated by antioxidant systems, as uncontrolled ROS can lead to oxidative stress (Apel and Hirt, 2004; Foyer and Noctor, 2009, 2011*b*; Trippi *et al*., 2009). Consequently, ROS are recognized as critical signaling molecules, and the cellular redox state is considered an intensive property of the living system that integrates information from metabolic, energetic and environmental cues controlling growth, development and acclimation responses, as well as cell suicide events (Breusegem and Dat, 2006; Robert *et al*., 2009, 2018; Trippi *et al*., 2009; Fernandez-göbel *et al*., 2019; Fichman and Mittler, 2020; Peláez-vico *et al*., 2022). In plants, chloroplast, mitochondria and peroxisome, which are redox active organelles where carbon and energetic metabolisms take place, are major sources of ROS production (Apel and Hirt, 2004). In chloroplasts, the photosynthetic process involves physical, biophysical, and biochemical redox reactions with different reaction rates. In consequence, oxygen became a potent univalent electron or energy acceptor from highly activated intermediates under conditions of excess excitation energy (EEE) induced by the different stress conditions (Karpinski *et al*., 1999; Foyer and Hanke, 2022). ROS produced in the chloroplasts are particularly important in the retrograde signaling of EEE conditions, affecting nuclear gene expression, acclimation responses as well as programmed cell death (Karpinski *et al*., 1999; Breusegem *et al*., 2001; Breusegem and Dat, 2006; Bechtold *et al*., 2008; Fichman and Mittler, 2020; Foyer and Hanke, 2022).

Likewise, several stress conditions in plants have been also shown to trigger or increase autophagy (Signorelli *et al*., 2019; Wang *et al*., 2021). Autophagy is an ancient and conserved process in eukaryotes by which different cellular components are delivered to vacuole/lysosome to be degraded, and is essential for plant energetic metabolism, growth, development and responses to different stress conditions (Su *et al*., 2020; Wang *et al*., 2021). Different types of autophagy have been identified in plants (Yagyu and Yoshimoto, 2023), being macroautophagy (hereafter termed autophagy) the most thoroughly studied. The autophagic process is initiated, and mainly modulated in the cytosol, by the formation of a double membrane cup, the phagophore, that expands to engulf macromolecules and cellular components to finally form a double-membrane vesicle called autophagosome. Autophagosomes are then transported and fused with the vacuole, facilitating the delivery of their cargo for subsequent degradation (Yagyu and Yoshimoto, 2023). The sequential steps of the autophagic process are coordinated by ATG (AuTophaGy) proteins as well as by some components of the endosome pathway, as recently reported (Zhuang *et al*., 2015; Robert *et al*., 2021). While early evidence pointed to a strong transcriptionally regulation of autophagy in response to nutritional starvation conditions and foliar senescence (Yoshimoto *et al*., 2004; Breeze *et al*., 2011), recent studies have shed light on the widespread modulation of autophagy initiation and progression by post-translational modifications of core autophagy proteins. These modifications primarily occur at the cytoplasmic level in response to environmental cues (Feng *et al*., 2015; Yang *et al*., 2019; Qi *et al*., 2021; Agbemafle *et al*., 2023).

It has been indicated a positive correlation between ROS generation and the autophagic process, where ROS can serve as both sources of damage and signals (Xiong *et al*., 2007*a*,*b*; Signorelli *et al*., 2019). Autophagy promotes the degradation of ROS-overproducing damaged organelles including chloroplasts, mitochondria, and peroxisomes, thus fulfilling antioxidant roles (Xiong *et al*., 2007*b*; Shibata *et al*., 2013; Nakamura *et al*., 2018; Oikawa *et al*., 2022). Supporting this notion, observations in different autophagy-mutant plants demonstrate the accumulation of oxidized and ubiquitinated proteins (Xiong *et al*., 2007*a*; Munch *et al*., 2014), high levels of H_2_O_2_ (Yoshimoto *et al*., 2009), increased amounts of total glutathione (Masclaux-Daubresse *et al*., 2014), accelerated senescence (Doelling *et al*., 2002; Hanaoka *et al*., 2002; Yoshimoto *et al*., 2004), and hypersensitivity to different stress conditions including oxidative stresses (Xiong *et al*., 2007*b*; Liu *et al*., 2009; Zhou *et al*., 2013), indicating a significant redox imbalance. On the other hand, oxidative treatments with H_2_O_2_ and methyl viologen (MV) have been shown to induce autophagy (Xiong *et al*., 2007*b*; Pérez-Pérez *et al*., 2012*b*). More recently, it has been hypothesized that H_2_O_2_ accumulation and membrane damage may mediate chlorophagy induced after UV-B and high light treatments in Arabidopsis leaves (Izumi *et al*., 2017; Woodson, 2019). Consequently, it is plausible that autophagy induction occurs as a response to the accumulation of cellular components that have been damaged. Notably, recent studies conducted in yeast, animals and Chlamydomonas have revealed that certain proteins involved in the autophagic process can be inhibited by ROS in *in vitro* experiments or artificial oxidative stress *in vivo*, indicating that ROS might suppress the activity of some autophagy-related proteins (Pérez-Pérez *et al*., 2014, 2016; Frudd *et al*., 2018; Mallén-ponce and Pérez-pérez, 2023). These findings raise apparently opposite evidence regarding the autophagy regulation by ROS. In this sense, a key aspect to be considered is the differential effects of ROS on autophagy, which varies both during oxidative stress and after the imposition of oxidative stress. Hence, the picture about the relationship among different ROS, subcellular redox changes, and autophagy regulation in plants is still unclear. Under this scenario, our study aimed to address the effects of different ROS on autophagy in plants, focusing on both the stress period and the post-stress recovery phase.

Herein, we investigated the effects of different ROS on autophagy in Arabidopsis seedlings. We demonstrate that oxidative treatments with H_2_O_2_ and MV, which drastically increase cytoplasmic oxidation, suppress autophagic flux, while restoration to a reduced cytoplasmic state -such as after stress subsides-correlates with the activation of autophagy. In contrast, chloroplast-generated ^1^O_2_ enhances autophagy without inducing statistically significant changes in cytoplasmic redox state. These findings provide novel insights into the complex interplay between ROS and autophagy, emphasizing the distinct roles of specific ROS in regulating this critical stress response. In summary, our findings unveil the distinct effects of ROS on autophagic flux, highlighting a strong correlation between autophagy and the physicochemical properties of the cytoplasm. Autophagy is inhibited under highly oxidized cytoplasmic conditions but is induced during recovery, likely playing a crucial role in removing damaged components.

## MATERIALS AND METODS

### Plant materials and growth conditions

*Arabidopsis thaliana* ecotype Columbia (Col-0) was used in this study. 35S:GFP-ATG8a transgenic seeds (CS39996) (Thompson et al., 2005) were obtained from the Arabidopsis Biological Resource Center. The cytosol-targeted fluorescent redox sensors GFP2-Orp1 or GRX-roGFP2 were provided by Dr. Markus Schwarzländer. Fluorescent (*flu*) and ferrochelatase 2 (*fc2*) mutants were provided by Dr. Jesse Woodson. We introgressed the GFP-ATG8a, GFP2-Orp1 and GRX-roGFP2 transgenes into the *flu* and *fc2* mutant lines by genetic crosses. Homozygous offspring were then selected from the F3 backcross generations.

Seeds were surface-sterilized and sown on culture plates containing nutrient-rich Murashige and Skoog (MS) solid medium lacking sucrose and containing 1 % agar. After sowing, plates were kept in darkness at 4°C for 2 to 3 days, and then transferred to a growth chamber under a 16 h light/8 h dark photoperiod. Photosynthetic photon flux density was 110 µmol m^-2^ sec^-1^. Average temperature was 22°C. Plants were grown for 7 days under these conditions and then passed to MS liquid medium to carry out the different treatments. For salt and osmotic stresses, NaCl (50 mM) and mannitol (100 mM) were used, respectively. Oxidative treatments were performed by the supplementation of the MS liquid medium with 1 µM methyl viologen (MV), 10 mM H_2_O_2_, 20 µM 3-(3,4-dichlorophenyl)-1,1-dimethylurea (DCMU), 20 µM methylene blue (MB). To scavenge superoxide and peroxide, 10 mM tiron and 10 mM benzoic acid (BA) were co-administrated. 10 mM dithiothreitol (DTT) was used as a reducing agent for protein disulfide bonds.

### Confocal fluorescence microscopy and image analysis

Confocal microscopy was performed on an Inverted microscope Nikon Eclipse CZ1. 35S:GFP-ATG8a expressing plants were observed with a 488-nm argon laser (BP 495–530 nm). To analyze autophagic bodies accumulation, 1 µM Concanamycin A was added to the different treatments. The number of autophagic bodies in epidermal cells was quantified after 16 h of treatments using ImageJ by counting their numbers in the central vacuole of multiple cells in sections of 50000 µm^2^. A macro for automating autophagic bodies counting was developed in ImageJ. Briefly, the input images were converted into an 8-bit format, followed by background subtraction using a rolling ball algorithm with a 50-pixel radius. Subsequently, particle analysis was performed on the binary image, considering particle size, circularity, and displaying the resulting masks.

Cytosol-targeted fluorescent redox sensors GFP2-Orp1 or GRX-roGFP2 were excited at 488 nm and 405 nm and fluorescent emissions were collected at 505-535 nm. These transgenic plants express a modified GFP sensitive to redox state (roGFP) genetically fused with different redox-active proteins, the peroxidase Orp1 and Glutaredoxin GRX1, which facilitate the sensing of H_2_O_2_ and the GSH ratio, respectively (Schwarzländer *et al*., 2009, 2015; Nietzel *et al*., 2019). The relative intensity of the two fluorescence excitation maxima depends on the redox state of roGFP, allowing ratiometric measurements (ratio 405/488) independent of probe concentration (Schwarzländer *et al*., 2015; Nietzel *et al*., 2019). roGFP predominantly equilibrates with the GSSG/2GSH couple, dependent on glutaredoxins (GRX). The transcriptional fusion of roGFP with GRX1 (GRX1-roGFP2) allows comparisons between different cell types and subcellular locations, as it is independent of endogenous GRX availability and insensitive to a wide pH range (pH 5.5-8.5) (Schwarzländer *et al*., 2015). Thiol peroxidases (Orp1) react extremely sensitively with H_2_O_2_, forming a sulfenic acid group and subsequently a disulfide bond. The oxidation of either of these groups can then be transferred, via a thiol-disulfide exchange reaction, to reductive systems such as thioredoxins or GRX, or to specific target proteins. The fusion of roGFP2 with Orp1 ensures that oxidation is efficiently passed from Orp1 to roGFP2. Dynamic measurements can be made because, once oxidized, roGFP2-Orp1 can be reduced again by redox couples (GRX and/or Thioredoxins (TRX) systems) (Schwarzländer *et al*., 2015; Nietzel *et al*., 2019). Therefore, the redox state of roGFP2-Orp1 is influenced not only by H_2_O_2_ but also by reducing agents (GSH and TRX). While the integration of oxidizing (H_2_O_2_) and reducing (GSH/TRX) influences might be considered a disadvantage of the roGFP2-Orp1 probe, they reflect how the redox state of endogenous thiols reactive with H_2_O_2_ is regulated, including the reducing capacity of the system. All imaging parameters were kept constant throughout all measurements. Subsequently, a single ratiometric image was generated using the ImageJ software, where the value of each pixel is the product of the ratio between the intensity at 405 nm with respect to that obtained at 488 nm according to (Schwarzlander *et al*., 2008). Quantitative measurements were calculated as the ratio of the mean intensity from each channel. The dynamic range of these reporter lines was established using 10 mM DTT and a curve of increasing concentrations of H_2_O_2_. For pseudocolor display, the ratio was coded on a spectral color scale ranging from blue (fully reduced) to red (fully oxidized).

### Immunoblot analyses

For GFP-ATG8 cleavage assay, seven-day old 35S:GFP-ATG8a transgenic plants were transferred to MS liquid medium to carry out the different treatments for 6 to 8 h. Then, total proteins were extracted. The extracts were centrifuged at 10 000 g for 10 min, and equal amounts of proteins from the supernatants were subjected to SDS-PAGE and immunoblotted with anti-GFP antibodies (Roche). Comparisons of absolute integrated density values for each line (GFP-ATG8 and free GFP) were made. Corresponding background values were subtracted from integrated density values, then integrated density values for bands corresponding to free GFP were expressed as relative values of GFP-ATG8 present in the same sample, and relativized to a control condition.

### Physiological parameters

Seven-day old Arabidopsis seedlings were exposed to the different treatments for 8 h and then the production of H_2_O_2_ and O_2_ ^-^, quantum efficiency of PSII photochemistry (ΦPSII), as well as the content of chlorophyll, soluble sugars and malondialdehyde (MDA) were analyzed. To assess H_2_O_2_ and O_2_ ^-^ accumulation, leaves were stained with DAB and NBT, respectively. The staining was quantified by measuring the intensity per area using ImageJ. Quantum efficiency of PSII photochemistry (ΦPSII) was measured from the upper surface using a pulse amplitude modulated fluorometer (FMS2, Hansatech Instruments, Pentney King’s Lynn, UK). Measurements were performed 6 h after the growth chamber lights were turned on. For chlorophyll, soluble sugars, and MDA analysis an alcoholic extract (1:2 w/v) was obtained from the samples. After centrifugation for 15 minutes at 12,000 *g*, the supernatant was diluted in ethanol, and the absorbance was measured at 654 nm using a spectrophotometer. Total chlorophylls were calculated following the method reported by (Wintermans M. and De Mots A., 1965). For soluble sugars determination, an aliquot of the sample was dried, and resuspended in a same volume of water. Then, the concentration of sugars was measured with anthrone (Fales, 1951) evaluating the absorbance at 620 nm using a glucose standard curve. For MDA determination, a volume of the supernatant was mixed with a same volume of a mixture of 0,5 % TBA and 20 % TCA, and incubated 20 minutes at 90 °C. The mixture was centrifuged at 12,000 *g* for 5 minutes, and the absorbance was measured at 532 and 600 nm (Heath and Packer, 1968).

### RNA extraction and RT-qPCR

Samples were homogenized in a cold mortar with TRIzol Reagent, mixed for 1 min and incubated at room temperature for 5 min. Then, 0.2 ml chloroform per ml of TRIzol Reagent was added and incubated at room temperature for 3 min. After incubation, the samples were centrifuged at 12 000 *g* at 4 °C for 15 min and the aqueous phases were transferred to clean tubes. RNA was precipitated by adding 1 vol of isopropanol, incubated at room temperature for 10 min and centrifuged at 12 000 *g* at 4 °C for 15 min. The precipitate was washed with 70 % ethanol and the samples were centrifuged again at 12 000 *g* at 4 °C for 15 min. The precipitates were dried and resuspended in diethyl pyrocarbonate-treated water and its concentration was quantified using a NanoDrop 3300 spectrometer (Thermo Scientific). Purified RNA was treated with DNase I (Invitrogen) to remove genomic DNA, according to the manufacturer’s instructions. DNA-free RNA (1 to 2.5 μg) was used with oligo(dT) for first strand cDNA synthesis using the Moloney murine leukemia virus reverse transcriptase (M-MLV RT, Promega) according to the manufacturer’s instructions. For each primer pair, the presence of a unique product of the expected size was checked on ethidium bromide-stained agarose gels after PCR reactions. Absence of contaminant genomic DNA was confirmed in reactions with DNase-treated RNA as template. The qPCR reaction was performed using iQ Universal SYBR Green Supermix (Bio-Rad). Amplification of actin with the forward primer 5’-TTCAGATGCCCAGAAGTCTTG-3’ and the reverse primer 5-GCGATACCTGAGAACATAGTGG-3 was used to normalize the amount of template cDNA. The specific primers pair employed for the detection of *Atg6* was forward primer 5’-AAAACTGGGAGAGCGATGAG-3’ and reverse primer 5’-GATAGATGAGCCCTGCGTAG-3’, for *Atg4a* was 5’-CCTTTGGAATCACTCGACCCATCT-3’ and 5’-ATCCGCAAACCCGTAGTTGCTT-3’, for *Atg7* was 5’-GCGGCTTTAGGTTTTGACAG-3’ and 5’-GTTGGTCTAGAGTTCGATCAGTC -3’, and for *Atg5* was 5’-TCCTGTTCGGTTGTATGTTCG-3’ and 5’-CGCTCTGTCTCCCATAAACTC-3’. qPCR was performed in thermocycler iQ5 (Bio-Rad) at 59 °C with iQ SYBR Green Supermix (Bio-Rad), according to the manufacturer’s instruction. Relative transcript abundance was calculated (Livak and Schmittgen, 2001).

### Microarray analysis

Publicly available microarray data were utilized to compare the expression of *Atg* genes under various conditions, including nitrogen deficiency (Rubin *et al*., 2009), treatment with 10 mM H_2_O_2_ for 24 h (Willems *et al*., 2016), treatment with 10 μM MV for 2 h (Scarpeci *et al*., 2008), and the examination of *flu* mutant plants subjected to an 8-hour cycle of darkness followed by 2 h of light (Op Den Camp *et al*., 2003). Expression values were normalized to the corresponding control conditions employed in each study and were expressed as log_2_ values.

### Statistical analyses

Data were analysed by analysis of variance (ANOVA) followed by the Di Rienzo–Guzman– Casanoves (DGC) test model, a cluster-based method for identifying groups of non-homogeneous means, using InfoStat software (Di Rienzo *et al*., 2002).

## RESULTS

### H_2_O_2_ and methyl viologen treatments induced cytoplasmic oxidation and inhibited autophagy in Arabidopsis

Initially, to gain insights into the interplay between ROS, redox state changes and autophagy, *Arabidopsis thaliana* seedlings were subjected to well-established autophagy-inducing conditions, namely nutritional starvation and different abiotic stress treatments (Wang *et al*., 2021), and both autophagy and the cytoplasmic redox state were analyzed (Fig. 1). To monitor autophagy, 35S:GFP-ATG8a (GFP-ATG8) transgenic plants were grown 7 days in MS solid medium, and then transferred to the different treatments in MS liquid medium supplemented with 1 µM Concanamycin A (Conc A) for a 16 hour-period. Conc A is a V-ATPase inhibitor widely used to prevent vacuole acidification, thereby effectively inhibiting the lytic activity of vacuoles, thus allowing the detection of autophagic bodies accumulation into the vacuoles through confocal microscopy (Fig. 1A and B). Analysis of autophagic flux using Conc A better defines autophagic activity since the increase of autophagosome number in the cytoplasm does not necessarily corresponds to an enhanced autophagic activity (Robert *et al*., 2021). When subjected to carbon and nitrogen starvation treatments, epidermal cells showed a high accumulation of autophagic bodies in the vacuole compared to control conditions, indicating an activated autophagic flux (Fig. 1A and B). Similarly, saline and osmotic stress treatments resulted in comparable outcomes, with a significant increase in autophagic bodies accumulating within the vacuoles (Fig. 1A and B). Besides, to evaluate the cytoplasmic redox state *in vivo*, 7 days old transgenic plants expressing the cytosol-targeted fluorescent redox sensors GFP2-Orp1 or GRX-roGFP2 were subjected to the different treatments (Fig. 1). These transgenic plants express a redox-sensitive green fluorescent protein (roGFPs) genetically fused to different redox-active proteins, responsive to H_2_O_2_ production and GSH:GSSG, respectively (Schwarzländer *et al*., 2009; Aller *et al*., 2013; Schwarzla *et al*., 2015; Nietzel *et al*., 2019). The relative intensity of the two fluorescence excitation maxima depends on the redox state of the modified GFP, enabling ratiometric measurements, independent of the probe concentration (Schwarzländer *et al*., 2015) (more details in *Materials and Methods*). The dynamic range of these reporter lines was previously monitored using DTT and H_2_O_2_ (Fig. S1A). Notably, in epidermal cells of seedlings expressing either GFP2-Orp1 or GRX-roGFP2, no changes in the cytoplasmic redox state were detected after 16 h of nutritional starvation, neither under saline or osmotic stress treatments compared to control conditions (Fig. 1A and C). These findings align with earlier and recent assessments of the redox state in cells of GRX-roGFP2 and roGFP2-ORP1 transgenic plants under water deficit and saline stress conditions (Schwarzländer *et al*., 2009; Bohle *et al*., 2024).

**Figure 1:**
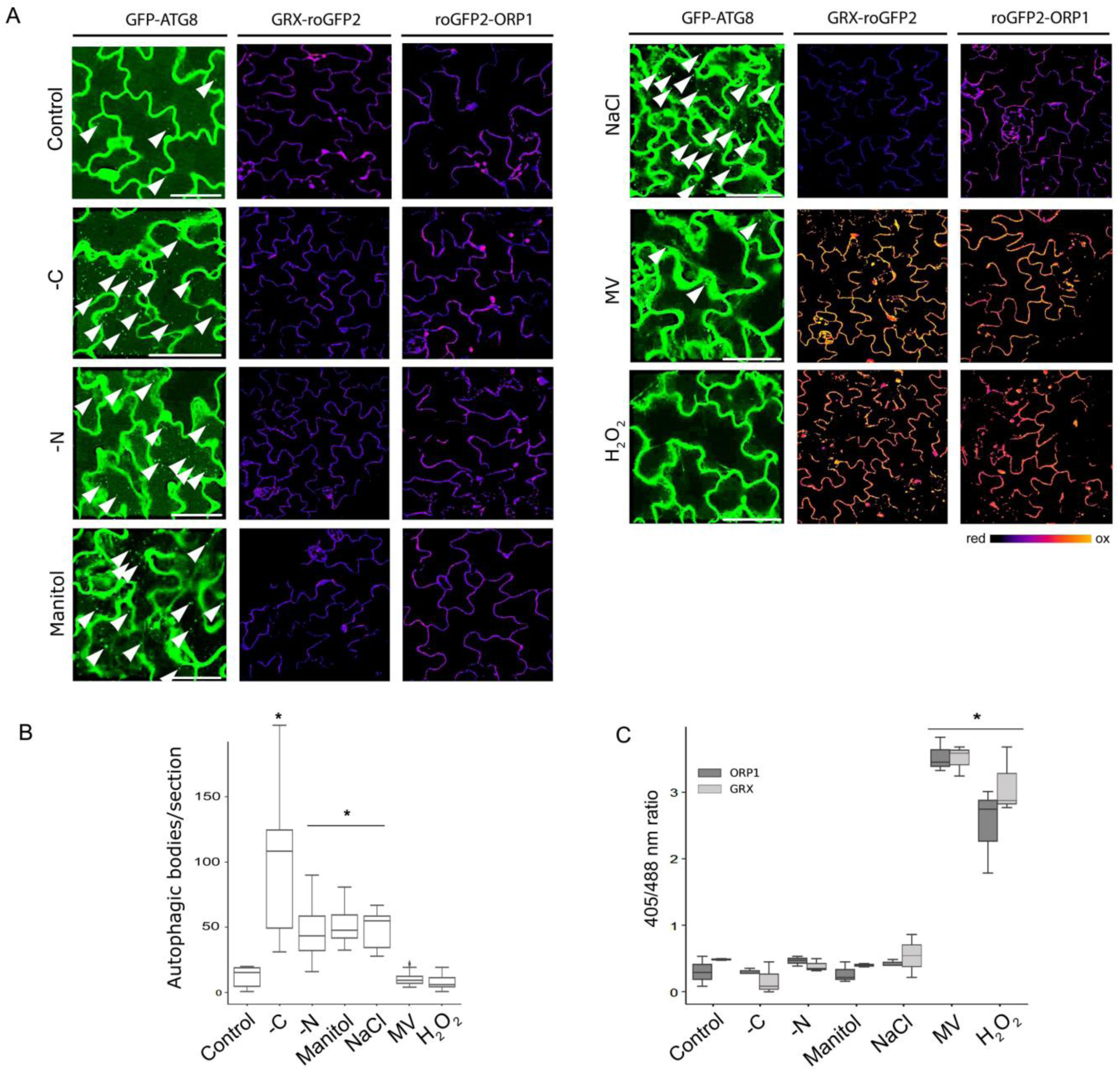
Autophagic flux and cytoplasmic redox state in *Arabidopsis thaliana* plants exposed to different abiotic stress conditions. (A-C) Seven-day old 35S:GFP-ATG8a transgenic plants and cytosol-targeted fluorescent redox sensors GFP2-Orp1 and GRX-roGFP2 expressing plants were exposed to different stress conditions in MS liquid medium. To induce autophagic bodies accumulation, MS liquid medium was supplemented with 1 μM Concanamycin A. (A) Representative images showing autophagic bodies accumulation and cytoplasmic redox state. The stress treatments were performed for 16 h, followed by the evaluation of autophagic bodies accumulation (GFP-ATG8 columns) and the cytoplasmic redox state (GRX-roGFP and GFP2-Orp1 columns) by confocal microscopy. Arrowheads indicate some of the autophagic bodies observed. Scale bars: 30 μm. GRX-roGFP and GFP2-Orp1 columns are ratiometric images. For pseudocolor display, the ratio was coded on a spectral color scale ranging from blue (fully reduced) to red (fully oxidized). The treatments included: MS with nitrogen (control), MS with nitrogen in darkness (-C), MS without nitrogen (-N), MS with nitrogen supplemented with 100 mM mannitol (Manitol), 50 mM NaCl (NaCl), 1 μM methyl viologen (MV) and 10 mM H_2_O_2_. (B) Quantification of autophagic body accumulation was performed using single-plane confocal images. Two sections from at least 6 plants were analyzed. (C) Quantification of cytoplasmic redox state expressed as the ratio of the fluorescence intensities measured at 488 nm and 405 nm. At least 4 plants were analyzed. Statistically significant differences are indicated by asterisks (\**P* < 0.05; error bars: means ± SEM, DGC test).

To further investigate the interplay between ROS, redox state changes and autophagy, oxidative stress treatments using H_2_O_2_ and methyl viologen (MV) were conducted. We evaluated different concentrations of H_2_O_2_ and observed a dose-dependent response for both the cytoplasmic redox state and autophagy inhibition. A similar approach was taken with MV, an herbicide widely used for the induction of photooxidative stress as it takes electrons from photosystem I, generating O_2-_ and H_2_O_2_ inside chloroplasts in the presence of light (Mehler, 1951; Farrington *et al*., 1973; Lascano *et al*., 2012; Nazish *et al*., 2022) (Fig. S1B-D). Based on these results and with the intention of employing moderate oxidative stress conditions that do not induce cell death, we used 10 mM H_2_O_2_ and 1 μM MV for subsequent experiments. Although oxidative treatments have been reported as autophagy-inducing conditions in Arabidopsis and Chlamydomonas (Xiong *et al*., 2007*b*; Pérez-Pérez *et al*., 2012*b*), no significant differences were observed in the accumulation of autophagic bodies in the epidermal cells of seedlings treated with MV or H_2_O_2_ for 16 h compared to the control condition (Fig. 1A and B). In contrast to observations under autophagy-inducing conditions such as nutritional starvation, saline, or osmotic stresses, these oxidative treatments significantly oxidized the cytoplasmic redox state, as revealed by analyzing both GRX-roGFP2 and roGFP2-ORP1 plants, as well as DAB staining, a histochemical reagent for H_2_O_2_ (Fig. 1A and C; Fig. S1; Fig. S2) (Fernandez-göbel *et al*., 2019).

To further characterize the effects of oxidative stress on autophagy, treatments with MV and H_2_O_2_ were performed on 35S:GFP-ATG8a plants under autophagic inducing conditions, specifically nitrogen and carbon starvation (Fig. 2, Fig. S2). Interestingly, the addition of 1 μM MV or 10 mM H_2_O_2_ during the 16-hour starvation treatments significantly reduced the accumulation of autophagic bodies (Fig. 2A and B). Furthermore, the inhibitory effects of the oxidative treatments on the autophagic flux were also confirmed by analyzing the degradation of the GFP-ATG8a fusion protein by western blot. This analysis provides insight beyond the cellular level, extending to the organ or plant level (Fig. 2D, Fig. S2). Furthermore, the evaluation of autophagic flux through the analysis of the degradation of the GFP-ATG8 fusion protein by immunoblotting allowed us to examine the effects of oxidative stress on autophagy in shorter time periods (6-8 h, Fig. 2D). Next, tiron and benzoic acid were co-administered with the MV treatment to scavenge superoxide and hydrogen peroxide, respectively (Navabpour *et al*., 2003; Mansouri *et al*., 2005; Velika and Kron, 2012; Liubimovskii *et al*., 2021). The effects of MV on autophagic flux, H_2_O_2_ accumulation, and cytoplasmic redox state were blocked in the presence of ROS scavengers (Fig. 2, Fig. S2A and B). Similarly, photooxidative stress induced by high light treatments showed an inhibitory effect on the autophagy flux, which was reversed in the presence of the superoxide radical scavenger, tiron (Fig. 2D). Moreover, both tiron and benzoic acid prevented photosynthetic damage caused by MV treatments, as measured by the quantum yield of photosystem II (ΦPSII), and prevented the decrease in soluble sugar levels (Fig. S2C). Furthermore, the inhibitory effects of MV on autophagic flux and the oxidation of the GSH:GSSG sensor probe (GRX-roGFP2) were reversed when 10 mM dithiothreitol (DTT), a reducing agent for protein disulfide bonds, was added (Fig. S2A and B). These results emphasize that the effects of MV in plants are associated with its capacity to induce ROS generation, contributing to a more oxidized cellular redox state and oxidative damage (Lascano *et al*., 2012). Notably, the restoration of autophagic flux in MV-treated plants by tiron, benzoic acid, and DTT suggests that H_2_O_2_ may inhibit autophagy, potentially by oxidizing disulfide bonds in crucial ATG proteins (Fig. S3 and S4) as demonstrated in recent studies in yeast, Chlamydomonas, and animals (Pérez-Pérez *et al*., 2014, 2016; Frudd *et al*., 2018; Zheng *et al*., 2020; Mallén-ponce and Pérez-pérez, 2023). H_2_O_2_ treatments inhibited autophagy in plant seedlings, roots, shoots, and detached mature leaves under carbon or nitrogen starvation conditions (Fig. S2D). Although the intensity of this effect may vary depending on the organ and developmental stage, probably due to differences in baseline redox states and cellular contexts for stress responses, these observations suggest a robust inhibitory role of H_2_O_2_ across various plant tissues.

**Figure 2:**
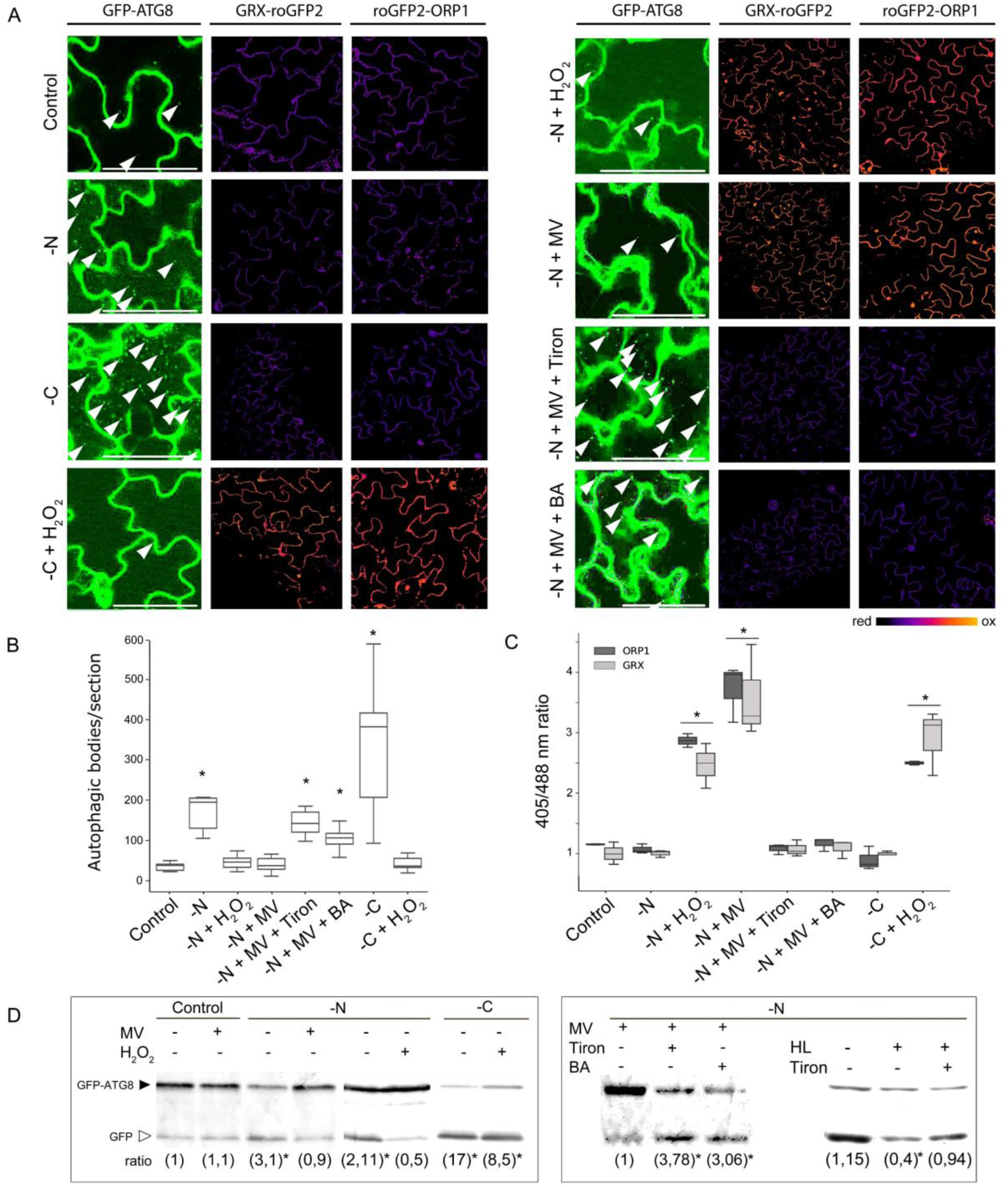
Treatments with MV and H_2_O_2_ highly oxidized the cytoplasm and prevented autophagy induction in Arabidopsis. (A-C) Seven-day old 35S:GFP-ATG8a transgenic plants and cytosol-targeted fluorescent redox sensors GFP2-Orp1 and GRX-roGFP2 expressing plants were exposed to autophagy-inducing conditions and the effect of methyl viologen (MV) and H_2_O_2_ on autophagic flux and cytoplasmic redox state were analyzed. (A) Representative images showing autophagic bodies accumulation and cytoplasmic redox state. The stress treatments were performed for 16 h, followed by the evaluation of autophagic bodies accumulation (GFP-ATG8 columns) and the cytoplasmic redox state (GRX-roGFP and GFP2-Orp1 columns) by confocal microscopy. To induced autophagic bodies accumulation, MS liquid medium was supplemented with 1 µM Concanamycin A. Arrowheads indicate some of the autophagic bodies observed. Scale bars: 30 μm. GRX-roGFP and GFP2-Orp1 columns are ratiometric images. The treatments included: MS with nitrogen (control), MS with nitrogen in darkness (-C) supplemented with 10 mM H_2_O_2_ (-C + H_2_O_2_), MS without nitrogen (-N) supplemented with 1 µM methyl viologen (-N + MV) or 10 mM H_2_O_2_ (-N + H_2_O_2_). 10 mM Tiron and 10 mM Benzoic acid (BA) were used for ROS scavenging (-N + MV + Tiron and -N + MV + BA). (B) Quantification of autophagic body accumulation was performed using single-plane confocal images. Two sections from at least 6 plants were analyzed. Two sections from at least 6 plants were analyzed. (C) Quantification of cytoplasmic redox state expressed as the ratio of the fluorescence intensities measured at 488 nm and 405 nm. At least 4 plants were analyzed. (D) Effects of oxidative treatments on the autophagic flux evaluated through the degradation of the GFP-ATG8a fusion protein by western blot. Seven-day old 35S:GFP-ATG8a transgenic Arabidopsis seedlings were exposed to the different treatments for 6-8 h. The GFP-ATG8a fusion has a predicted molecular weight of approximately 40 kDa (closed arrowheads); free GFP has a predicted molecular weight of 27 kDa (open arrowheads). Representative images are shown. Ratio: the numbers in brackets denote the GFP/GFP-ATG8 ratio normalized to the control condition (well 1 in each box). Three replicates were performed (more than fifteen plants per replicate). HL: high light treatment (≈1000 µmoles m^-2^ s^-1^). Asterisks indicate statistically significant differences (\**P* < 0.05; error bars: means ± SEM, DGC test).

### Autophagy is induced in the recovery phase after oxidative stress in Arabidopsis

Previous reports indicated that autophagy is involved in degrading oxidized proteins and organelles in response to oxidative stress (Xiong *et al*., 2007*b*; Shibata *et al*., 2013; Izumi *et al*., 2017; Oikawa *et al*., 2022). However, our experimental results demonstrate that oxidative stress treatments inhibit autophagy. Considering these findings, we aimed to investigate autophagic activity following oxidative stress treatments by assessing the autophagic flux in Arabidopsis plants during the recovery phase. With this purpose, 7-day-old Arabidopsis seedlings were subjected to a brief exposure of 10 mM H_2_O_2_ or high light intensity (≈1000 µmoles m^-2^ s^-1^) for 4 hours, followed by transfer to H_2_O_2_-free MS liquid medium under control conditions (≈100 µmoles m^-2^ s^-1^) for a 20-hour recovery period. Additionally, a set of plants were incubated in 10 mM H_2_O_2_ or to control conditions (H_2_O_2_-free MS liquid medium) for 24 h. This experimental design aimed to elucidate the dynamic changes in autophagy following oxidative stress and determine whether autophagic activity is modulated during the recovery phase. The results showed that autophagic flux was inhibited in plants incubated 24 h with H_2_O_2_ compared to control conditions (Fig. 3A, B and D). Interestingly, autophagic flux was not only recovered but also induced during the recovery period after the oxidative stress treatments surpassing the levels observed in plants subjected to control conditions (Fig. 3A, B and D). Likewise, after the 20-hour recovery period, the redox biosensors returned to their reduced state, indicating cytoplasmic redox recovery, and a well-defined correlation between the reduced cytoplasmic redox state and autophagic flux (Fig. 3A and C).

**Figure 3:**
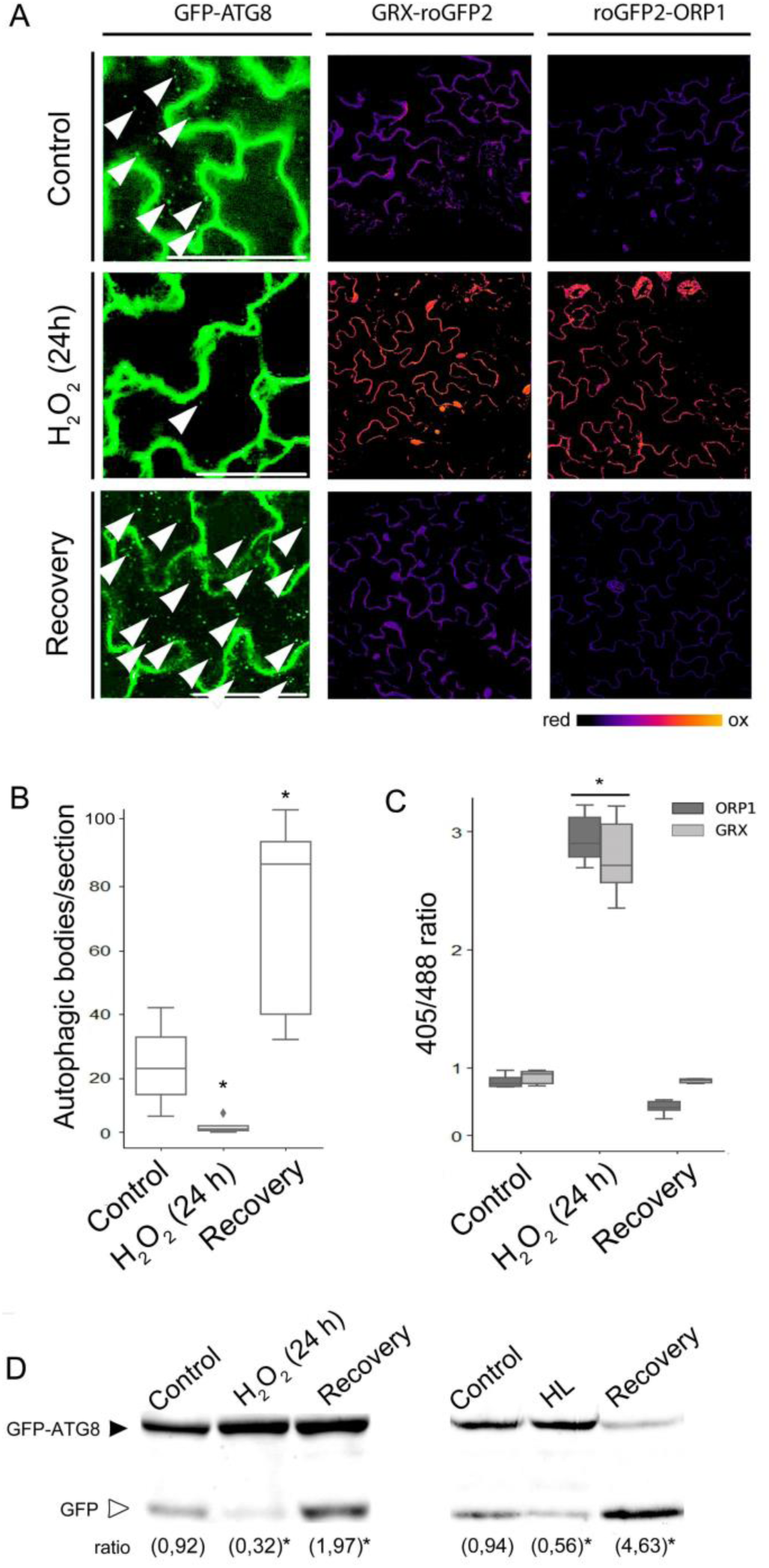
Autophagy is induced after oxidative stress treatments in Arabidopsis. (A-D) Seven-day old transgenic plants expressing 35S:GFP-ATG8a and cytosol-targeted fluorescent redox sensors GFP2-Orp1 and GRX-roGFP2 were subjected to a brief exposure of oxidative stress, either 10 mM H_2_O_2_ or high light intensity (≈1000 µmoles m^-2^ s^-1^), for 4 h. Subsequently they were transferred to MS liquid medium for a 20-hour recovery period under continuous light (≈120 µmoles m^-2^ s^-1^) (recovery). Control plants were incubated under continuous light (≈120 µmoles m^-2^ s^-1^) in H_2_O_2_-free MS liquid medium for 24 h. Stressed plants were incubated in MS supplemented with 10 mM H_2_O_2_ or high light intensity (≈1000 µmoles m^-2^ s^-1^) for 24 h. (A) Representative images showing autophagic bodies accumulation and cytoplasmic redox state. To induce accumulation of autophagic bodies, MS liquid medium was supplemented with 1 µM Concanamycin A. Arrowheads indicate some of the autophagic bodies observed. Scale bars: 30 μm. GRX-roGFP and GFP2-Orp1 columns represent ratiometric images, with pseudocolor display indicating the ratio on a spectral color scale ranging from blue (fully reduced) to red (fully oxidized). (B) Quantification of autophagic body accumulation was performed using single-plane confocal images. Two sections from at least 6 plants were analyzed. (C) Quantification of cytoplasmic redox state expressed as the ratio of the fluorescence intensities measured at 488 nm and 405 nm. At least 4 plants were analyzed. Asterisks indicate statistically significant differences (\**P* < 0.05; error bars: means ± SEM, DGC test). (D) Autophagic flux evaluated through the degradation of the GFP-ATG8a fusion protein by western blot. Total protein (40 µg) was subjected to immunoblot analysis with anti-GFP antibodies. The GFP-ATG8a fusion has a predicted molecular weight of approximately 40 kDa (closed arrowheads), while free GFP has a predicted molecular weight of 27 kDa (open arrowheads). Representative images are shown. The numbers in brackets denote the GFP/GFP-ATG8 ratio normalized to the control condition. Three replicates were performed (more than fifteen plants per replicate). Asterisks indicate statistically significant differences (\**P* < 0.05; error bars: means ± SEM, DGC test).

### Singlet oxygen induced autophagic flux without effect on the cytoplasmic redox state

Singlet oxygen (^1^O_2_) is mainly produced in the chloroplast, and its targets are predominantly confined inside this organelle due to the specific physicochemical characteristic of this ROS (Dmitrieva *et al*., 2020). To investigate the effects of ^1^O_2_ on the autophagic activity, GFP-ATG8a seedlings were treated with 3-(3,4-dichlorophenyl)-1,1-dimethylurea (DCMU) (Fig. 4), an herbicide that specifically induces ^1^O_2_ generation by blocking the electron transport chain in photosystem II and inhibiting the plastoquinone reduction. It has been reported that DCMU promotes ^1^O_2_ generation without affecting superoxide anion and hydroxyl radicals at PSII (Fufezan *et al*., 2002; Flors *et al*., 2006; Dmitrieva *et al*., 2020). The analysis of autophagic body accumulation and cytoplasmic redox state was conducted after a 16-hour treatment (Fig. 4A-C). Additionally, treatments with methylene blue (MB), a type-II photosensitizer able to generate ^1^O_2_ in leaves under visible light (Shao *et al*., 2007, 2013), with minor effects on photosynthesis (Fig. 4E), were also performed. Treatments with these chemicals increased the accumulation of autophagic bodies but did not show statistically significant effects on the redox state of the cytoplasm in Arabidopsis cells compared to control treatment (Fig. 4A and B). The autophagic flux-inducing effect of DCMU and MB was also evidenced after 6-8 hours of treatments through the analysis of GFP-ATG8 degradation by immunoblotting (Fig. 4D).

**Figure 4:**
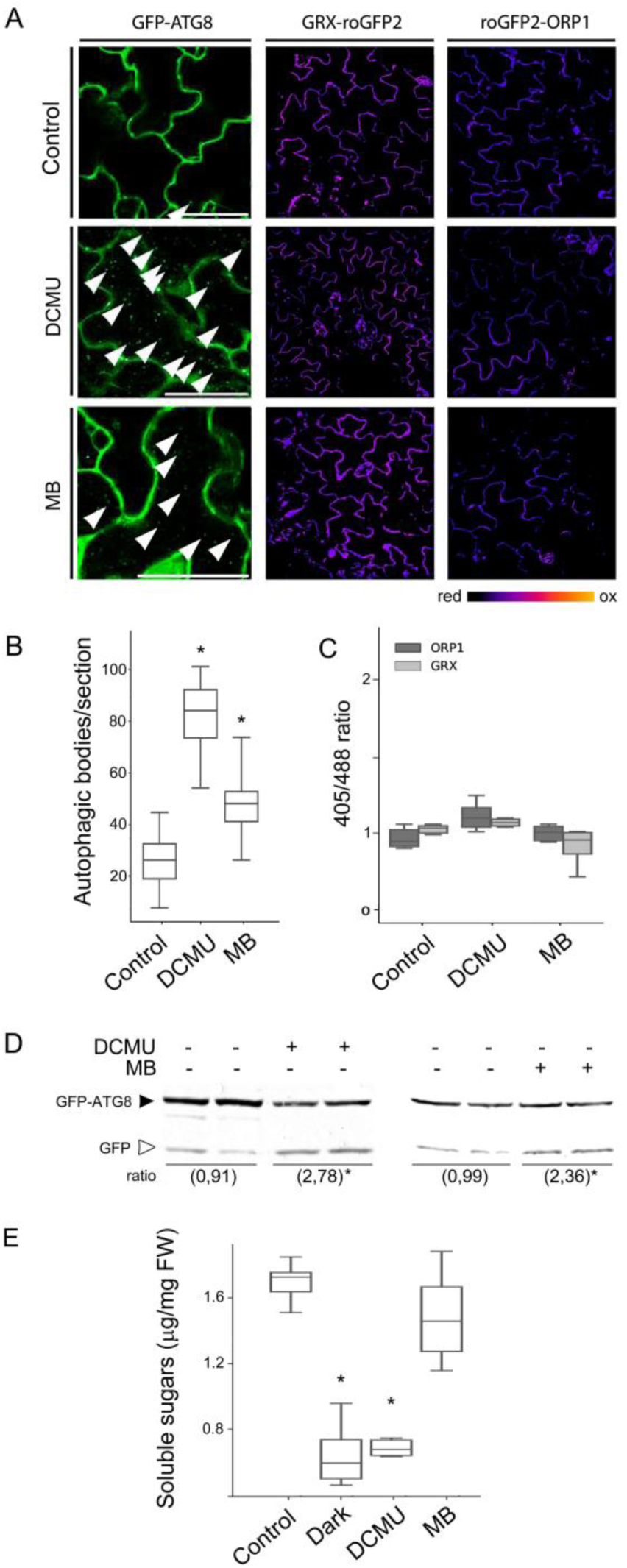
DCMU and methylene blue treatments induced autophagic flux in Arabidopsis. (A-D) Seven-day old transgenic plants expressing 35S:GFP-ATG8a and cytosol-targeted fluorescent redox sensors GFP2-Orp1 and GRX-roGFP2 were incubated in MS medium supplemented with either 20 μM 3-(3,4-dichlorophenyl)-1,1-dimethylurea (DCMU) or 20 μM methylene blue (MB), and the autophagic flux and cytoplasmic redox state were evaluated. (A) Representative images showing autophagic bodies accumulation and cytoplasmic redox state after 16 h of treatment. To induce autophagic bodies accumulation, MS liquid medium was supplemented with 1 μM Concanamycin A. Some of the autophagic bodies are indicated by arrowheads. Scale bars: 30 μm. The GRX-roGFP and GFP2-Orp1 columns represent ratiometric images. For pseudo color display, the ratio was coded using a spectral color scale ranging from blue (fully reduced) to red (fully oxidized). (B) Quantification of autophagic body accumulation was performed using single-plane confocal images. Two sections from at least 6 plants were analyzed. (C) Quantification of cytoplasmic redox state expressed as the ratio of the fluorescence intensities measured at 488 nm and 405 nm. Analysis was performed on at least 4 plants. Asterisks indicate statistically significant differences (\**P* < 0.05; error bars: means ± SEM, DGC test). (D) Evaluation of the effects of DCMU and MB on autophagic flux through degradation of the GFP-ATG8a fusion protein by western blot. Seven-day old transgenic Arabidopsis seedlings expressing 35S:GFP-ATG8a were exposed to the different treatments for 6-8 h. Total protein (40 μg) was subjected to immunoblot analysis with anti-GFP antibodies. The GFP-ATG8a fusion has a predicted molecular weight of approximately 40 kDa (closed arrowheads), while free GFP has a predicted molecular weight of 27 kDa (open arrowheads). Representative images are shown. The numbers in brackets denote the GFP/GFP-ATG8 ratio normalized to control condition. Four replicates were performed with more than fifteen plants per replicate. (E) Seven-day old Arabidopsis seedlings were exposed to the different treatments and soluble sugar content was analyzed. FW: fresh weight. Asterisks indicate statistically significant differences (\**P* < 0.05; error bars: means ± SEM, DGC test).

To further investigate the effects of ^1^O_2_ on the autophagic process, we performed experiments with widely used endogenously chloroplast ^1^O_2_-hyperproducing *Arabidopsis* mutant plants: fluorescent (*flu*) and two different lines of ferrochelatase 2 (*fc2*), *fc2.1* which presents reduced expression levels of *Fc2*, and the null mutant *fc2.2*. These mutant plants accumulate different photosensitizing intermediates of chlorophyll biosynthesis in the dark, which, upon light exposure lead to an exacerbated production of ^1^O_2_ (Op Den Camp *et al*., 2003). The *flu* mutant plants accumulate protochlorophyllide (Meskauskiene *et al*., 2001), while mutations in plastid ferrochelatase 2 lead to the accumulation of protoporphyrin IX (Scharfenberg *et al*., 2015). Likewise, these genotypes were crossed with autophagy reporter-and redox sensor-transgenic plants. Initially, wild type, *flu* and *fc2* mutant plants were grown 7 days under continuous light conditions (control conditions), and several growth and physiological parameters were evaluated (Fig. S6). Interestingly, the *fc2* mutants showed arrested root growth, reduced chlorophyll levels, increased levels of malondialdehyde (MDA), an intermediary metabolite of lipid peroxidation used as oxidative stress marker (Robert *et al*., 2014), and reduced levels of soluble sugars compared to wild type, while no significant differences were observed between wild type and *flu* mutant (Fig. S6). Likewise, the cytoplasmic redox state in *fc2* plants was significantly more oxidized than that in the wild type and *flu* backgrounds (Fig. S6, Fig. 5). Furthermore, and consistent with a previous study (Lemke *et al*., 2021), the autophagic flux was induced in *fc2* mutants plants compared to wild type and *flu* mutant under continuous light (control) conditions (Fig. S6). Hence, considering the altered phenotypes observed in *fc2* mutants even under control conditions, we used the *flu* mutants for subsequent analysis.

**Figure 5:**
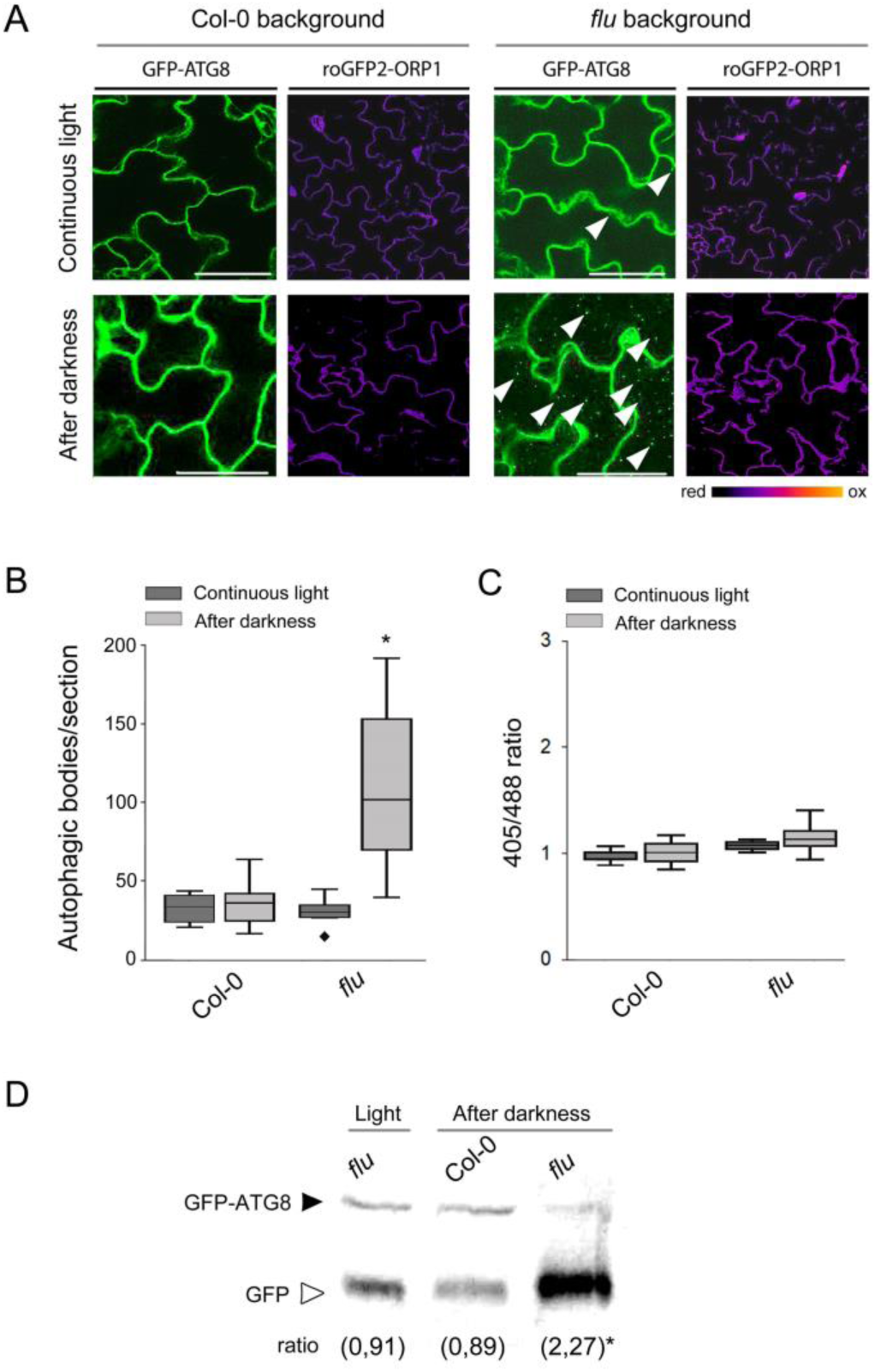
Autophagic flux is induced in conditionally chloroplast ^1^O_2_-hyperproducing *flu* mutant Arabidopsis plants. (A-C) Seven-day old wildtype (Col-0) and *flu* mutant plants expressing 35S:GFP-ATG8a or GFP2-Orp1 were transferred to darkness for 8 h, followed by re-exposure to light. Control plants were maintained under continuous light conditions. After 8 h, the autophagic flux and cytoplasmic redox state were analyzed. (A) Representative images showing autophagic bodies accumulation and cytoplasmic redox state. To induce autophagic bodies accumulation, the MS liquid medium was supplemented with 1 μM Concanamycin A. The GFP2-Orp1 columns represent ratiometric images. For pseudocolor display, the ratio was coded on a spectral color scale ranging from blue (fully reduced) to red (fully oxidized). Arrowheads indicate some of the autophagic bodies observed. Scale bars: 30 μm. (B) Quantification of autophagic body accumulation was performed using single-plane confocal images. Two sections from at least 6 plants were analyzed. (C) Quantification of cytoplasmic redox state expressed as the ratio of the fluorescence intensities measured at 488 nm and 405 nm. At least 4 plants were analyzed. Asterisks indicate statistically significant differences (\**P* < 0.05; error bars: means ± SEM, DGC test). (D) Evaluation of autophagic flux through the degradation of the GFP-ATG8a fusion protein by western blot. Seven-day old wildtype (Col-0) and *flu* mutant plants expressing 35S:GFP-ATG8a were transferred to darkness for 8 h and then re-exposed to light for 6 h (after darkness). Continuous light was used as a control (light). Total protein (40 μg) was subjected to immunoblot analysis with anti-GFP antibodies. The GFP-ATG8a fusion has a predicted molecular weight of approximately 40 kDa (closed arrowheads); while free GFP has a predicted molecular weight of 27 kDa (open arrowheads). Representative images are shown. The numbers in brackets denote the GFP/GFP-ATG8 ratio normalized to *flu* mutans under control condition. Three replicates were performed (more than fifteen plants per replicate). Asterisks indicate statistically significant differences.

*flu*/GFP-ATG8a plants were grown under continuous light conditions for 7 days, transferred to darkness for 8 h, and then re-exposed to light in the presence of Conc A for another 6-8 h. After the light/dark/light cycling conditions, a significantly increased autophagic flux was observed in *flu*/GFP-ATG8a compared to Col-0/GFP-ATG8 plants. However, no statistically significant differences were observed among these genotypes under continuous light conditions (Fig. 5A, B and D). Besides, the cytoplasmic redox state was monitored under control conditions as well as after a period of darkness by analyzing the fluorescent redox sensor roGFP2-ORP1 expressed in the wild type and *flu* backgrounds. Although a slight tendency towards a more oxidized state was observed in the cytoplasm of *flu* mutant plants once re-exposed to light, the differences were not statistically significant (Fig. 5C). Thus, in contrast to H_2_O_2_ and MV treatments, ^1^O_2_ increased the autophagic flux and did not oxidize the cytoplasmic redox state, indicating differential effects on the autophagic activity depending on the ROS nature, subcellular site of action, and their effects on the cytoplasmic redox state. Additionally, the transcript levels of *Atg4*, *Atg5*, *Atg6* and *Atg7* were analyzed by RTq-PCR in *flu* mutant after 8 h of light re-exposure. No statistically significant differences were observed in the expression level of the analyzed genes between wild type and *flu* mutant neither under control conditions nor after cycling conditions (Fig. S7).

## DISCUSSION

Plants, as photosynthetic, aerobic and sessile organisms, naturally produce reactive oxygen species (ROS) as byproducts of various metabolic processes, especially when exposed to environmental stresses. Due to their high reactivity, ROS play dual roles, acting as agents of damage and as signaling molecules. Consequently, ROS are universally implicated in plant stress symptomatology, and also play a pivotal role in modulating plant growth, development, and cell death (Apel and Hirt, 2004; Trippi *et al*., 2009; Baxter *et al*., 2014; Choudhury *et al*., 2017; Fichman and Mittler, 2020). Similarly, a variety of stress conditions in plants have been demonstrated to initiate or enhance autophagy (Bassham *et al*., 2006; Liu *et al*., 2009; Signorelli *et al*., 2019; Su *et al*., 2020). Moreover, defects in antioxidant proteins in Arabidopsis and Chlamydomonas contribute to uncontrolled ROS bursts and autophagy induction (Pérez-Pérez *et al*., 2012*a*; Shibata *et al*., 2013). While the concurrent induction of ROS production and autophagy activation suggests a positive correlation between these cellular processes, certain molecular mechanisms reported in animal cells, yeast, and Chlamydomonas, indicate that ROS may also negatively influence on autophagy (Pérez-Pérez *et al*., 2014, 2016; Frudd *et al*., 2018; Mallén-ponce and Pérez-pérez, 2023). However, direct evidence of ROS-mediated inhibition of autophagy in plants remains scarce. Additionally, it is important to consider whether autophagy is activated directly by ROS signaling or as a response to cellular damage induced by ROS, potentially creating a feedback loop between ROS production and autophagic activity. Furthermore, existing literature often treats ROS as a homogeneous group, overlooking the potential differences in how distinct types of ROS impact autophagy, and not consistently distinguishing their effects during stress imposition versus recovery phases in plants. Our findings point in that direction, highlighting the differential effects of ROS on autophagy, and suggesting that under specific stress conditions leading to a highly oxidized cytoplasmic state, the autophagic process may be suppressed. Additionally, the impact of ROS on autophagy is likely influenced by the developmental stage and the specific organ, as these factors determine both the baseline redox state and the cellular context for stress responses. Once the oxidative stress subsides and the cytoplasmic redox status is restored, autophagy is activated, potentially facilitating the removal of damaged cellular components.

The relationship between autophagy and ROS is intricate, and our findings revealed that different types of ROS, even when generated in the same organelle, can have different effects on the autophagic process. We observed that under conditions of nutritional starvation, which are canonical inducers of autophagy, oxidative treatments with MV and H_2_O_2_, that led to significant oxidation of the cytoplasmic redox state, had an inhibitory effect on the autophagic flux (Fig. 2). This implies that the comparatively limited autophagosome biogenesis rate under optimal growth conditions could potentially account for the absence of observable inhibitory effects of ROS on autophagy in prior *in vivo* plant studies (Fig. 1). Opposite to the inhibitory effects of H_2_O_2_ on the autophagic flux, the specific generation of ^1^O_2_ in the *flu* mutant plants, as well as treatments with DCMU and MB (Fufezan *et al*., 2002; Op Den Camp *et al*., 2003; Flors *et al*., 2006; Shao *et al*., 2007, 2013; Dmitrieva *et al*., 2020), increased the autophagic activity (Fig. 4 and Fig. 5). These findings are consistent with previous research demonstrating that treatments with norflurazon, an inhibitor of carotenoid biosynthesis, triggers autophagy in *Chlamydomonas reinhardtii* (Pérez-Pérez *et al*., 2012*a*, 2016). Similarly to what is observed in *flu* mutants (Op Den Camp *et al*., 2003), norflurazon treatments lead to perturbations in plastid homeostasis that increase the photosensitizing activity of tetrapyrroles, which can transfer the excitation energy to the oxygen molecule, thereby generating ^1^O_2_ (Kim and Apel, 2013). Singlet oxygen, carotenoid derivatives, and tetrapyrroles are principal components of the retrograde signaling pathway, a key process that regulates not only chloroplastic processes but also intersects with other cellular signaling events playing broader roles in plant function (Chan *et al*., 2016).

The distinct effects of ^1^O_2_ and H_2_O_2_ on autophagy are likely driven by their unique chemical properties, reactivities, and molecular targets. ^1^O_2_, being highly reactive and short-live, primarily acts near its generation site. Consequently, no significant redox changes were observed in the cytoplasm following DCMU and MB treatments or in *flu* mutants (Fig. 4 and 5). This ROS primarily predominantly reacts with unsaturated fatty acids, generating hydroperoxides that disrupt lipid membranes and potentially signal damage, being the major ROS involved in photooxidative damage (Triantaphylidès *et al*., 2008). Chloroplastic membrane damage, in particular, may trigger specific autophagy-activating signals (Izumi *et al*., 2017; Nakamura *et al*., 2018). Conversely, treatments with MV and H_2_O_2_, which reduced autophagic flux, significantly oxidized the cytoplasmic redox state (Fig. 1 and Fig. 2). These findings establish a clear correlation between the cytoplasmic redox status and autophagic flux. H_2_O_2_ is less reactive but more diffusible than ^1^O_2_, allowing it to oxidize thiol groups in cysteine residues of proteins throughout the cell, leading to structural and functional alterations. Supporting this, MV and H_2_O_2_ -induced oxidation of the cytoplasmic redox state, which could be reversed by ROS scavengers such as tiron or benzoic acid, as well as by the disulfide bond reductant DTT. These observations suggest that H_2_O_2_ not only inhibits autophagy but also implicates protein oxidation in this response. At the molecular level, the activity of key ATG proteins has been demonstrated to be subjected to redox regulation, being the proteins active when reduced and inactive when oxidized. Different cysteine residues have been identified as targets for redox regulation (Figs. S3 and S4). For example, the protease ATG4 which cleaves ATG8 for lipidation and delipidation with PE, is inhibited by H_2_O_2_ in *Saccharomyces cereivisiae* and *Chlamydomonas reinhardtii*. This inhibition involves Cys400 and Cys338 in Chlamydomonas and yeast, respectively (Pérez-Pérez et al., 2014, 2016). Reducing agents like DTT can restore ATG4 activity (Pérez-Pérez et al., 2014, 2016). Similarly, oxidation of ATG3 and ATG7, leads to the formation of an intermolecular disulfide-bound complex between Cys264 in ATG3 and Cys572 in ATG7, inhibiting ATG8 lipidation in animal cells and recently observed in Chlamydomonas (Frudd *et al*., 2018; Mallén-ponce and Pérez-pérez, 2023). The presence of these amino acid residues among ATG protein isoforms in yeast, human cells, Chlamydomonas and Arabidopsis (Figs. S3 and S4) suggests that similar redox regulation may also occur in plants. These conserved cysteines and others identified as potential redox candidates (Figure S4) are targets to analyze in future studies.

The complexity of the relationship between autophagy and ROS increases when considering that autophagy plays a critical role in degrading oxidized proteins and damaged organelles, which, in turn, are both products of oxidative stress and primary sources of ROS production. This highlights a feedback loop where the degradation of ROS-damaged organelles via autophagy helps mitigate further ROS generation, yet the oxidative damage to these organelles also drives the need for autophagic degradation. A crucial question in this context is to what extent autophagy activation under oxidative stress is directly regulated by ROS or arises as a secondary response to ROS-induced damage. ROS can directly influence autophagic process through redox modifications of key proteins involved in the autophagy machinery. Paradoxically, it has been shown that oxidation of ATG components from yeast, animals and Chlamydomonas exerts a negative regulatory effect on their activity (Pérez-Pérez *et al*., 2014, 2016; Frudd *et al*., 2018; Mallén-ponce and Pérez-pérez, 2023). In parallel, oxidative stress causes cellular damage, such as the oxidation of proteins, lipids, and nucleic acids, or the disruption of organellar integrity. These byproducts of oxidative damage may act as signals that trigger autophagy as part of a cellular clearance and repair mechanism (Izumi *et al*., 2017; Nakamura *et al*., 2018, 2021). Understanding the balance between these direct and indirect pathways is critical to dissect the precise role of ROS in regulating autophagy. Our assessment of autophagic flux aimed to dissect this relationship, shedding light on the role of autophagy in cellular adaptation to oxidative stress and its recovery. Previous research has shown that photodamage-inducing treatments, such as high-intensity or UV-B light irradiation, stimulate chloroplast degradation *via* autophagy, while antioxidants treatments inhibited autophagy induction, suggesting that ROS accumulation or the resulting damage plays a role in triggering this process (Izumi *et al*., 2017). In our experiments, we observed increased autophagic flux during the recovery period after oxidative stress exposure, concurrent with the restoration of a reduced redox state in the cytoplasm, indicating that the autophagy inhibition caused by O_2-_/H_2_O_2_ is transient (Fig. 3). Our findings also suggest that the induction of autophagy after oxidative stress subsides is associated with the clearance of damaged cytoplasmic components, as evidenced by treatments with the antioxidants tiron and benzoic acid (Fig. S2C). These treatments decreased ROS production and mitigated cell injury, supporting the idea that autophagy plays a critical role in resolving oxidative damage. Additionally, the transcript analysis of *ATG* genes supports the notion that the observed inhibitory redox regulation primarily operates at a post-transcriptional and/or post-translational level. This aligns with existing literature suggesting oxidative modifications negatively impact the ATG machinery’s functionality, although further studies are required to confirm and explore these dynamics. Overall, these findings underscore the dynamic nature of autophagy in response to oxidative stress and recovery, suggesting a complex regulatory mechanism in plants.

In summary, our work provides a novel perspective on the relationship between ROS and autophagy. The distinct effects of H_2_O_2_ and ^1^O_2_ on autophagy underscore that the term “ROS” encompasses a diverse group of molecules impacting on this process in fundamentally different ways. While the relationship between ROS and autophagy in plants has been the focus of numerous studies, many investigations have treated ROS as a homogeneous category. This approach, while valuable in establishing foundational insights, may overlook the distinct effects that specific ROS types exert on autophagic processes. Our findings align with previous studies suggesting that H_2_O_2_ and ^1^O_2_ have antagonistic effects on plant stress responses (Karpinska *et al*., 2000; Laloi *et al*., 2007; Triantaphylidès *et al*., 2008; Melchiorre *et al*., 2009; Sabater and Martín, 2013). The differential effects of these ROS on autophagic flux observed in this study are consistent with earlier reports demonstrating distinct impacts of H_2_O_2_ and ^1^O_2_ on the activity of ATG4, reflected in the levels of lipidated ATG8 after treatments with H_2_O_2_ or norflurazon (Pérez-Pérez *et al*., 2016). These results highlight the potential for specific ROS to exert unique regulatory roles in the modulation of autophagy. Recognizing these nuances provides a deeper understanding of the intricate interplay between ROS signaling and autophagic processes. Further research should employ more advanced methodologies, such as quantitative global analysis through mass spectrometry or electron paramagnetic resonance (EPR), to precisely measure ROS dynamics in plants subjected to abiotic stress. These approaches are essential to correlate the relative abundance of distinct ROS with autophagy modulation, ultimately shedding light on their roles in modulating autophagy in response to fluctuating environmental conditions.

## SUPPLEMENTARY DATA

Supplementary Figure 1: Optimization of methyl viologen (MV) and H_2_O_2_ concentrations, and calibration of the redox sensors.

Supplementary Figure 2: DTT, tiron and benzoic acid reversed the effects of MV treatments.

Supplementary Figure 3: Sequence alignment of ATG4 proteases from different organisms.

Supplementary Figure 4: Sequence alignment of ATG3 and ATG7 proteins from different organisms.

Supplementary Figure 5: Characterization of *fc2* and *flu* mutant plants under continuous light conditions.

Supplementary Figure 6: Transcript level of *Atg* genes in Arabidopsis plants subjected to different oxidative treatments.

## ACKNOWLEDGEMENTS

We thank Dr. Leandro Ortega, Dra. Virginia Lobatto and Dra. Anahí Yañez (Unidad de Doble Dependencia INTA-CONICET (UDEA)) for helpful support with the confocal microscopy and for technical support. We also thank Sistema Nacional de Microscopia of the Ministerio de Ciencia, tecnología e innovación Productiva (MINCyT), Argentina.

## AUTHOR CONTRIBUTION

GR and RL: conceptualization; GR and AE: investigation; GR, AE, LS and RL: formal analysis; GR, LS, and RL: resources; GR and AE: writing original draft and incorporated feedback from all authors; GR and AE: visualization; GR and RL: supervision; GR, LS and RL: funding acquisition.

## ABBREVIATIONS

ATG: autophagy genes
Conc A: Concanamicina A
GFP: green fluorescent protein
GSH: reduced glutathione
GSSG: oxidized glutathione
MV: methyl viologen
roGFP: redox-sensitive green fluorescent proteins
ROS: reactive oxygen species

