## Supplementary data for "Redox regulation of autophagy in Arabidopsis: differential effects of reactive oxygen species"

### SUPPLEMENTARY FIGURES

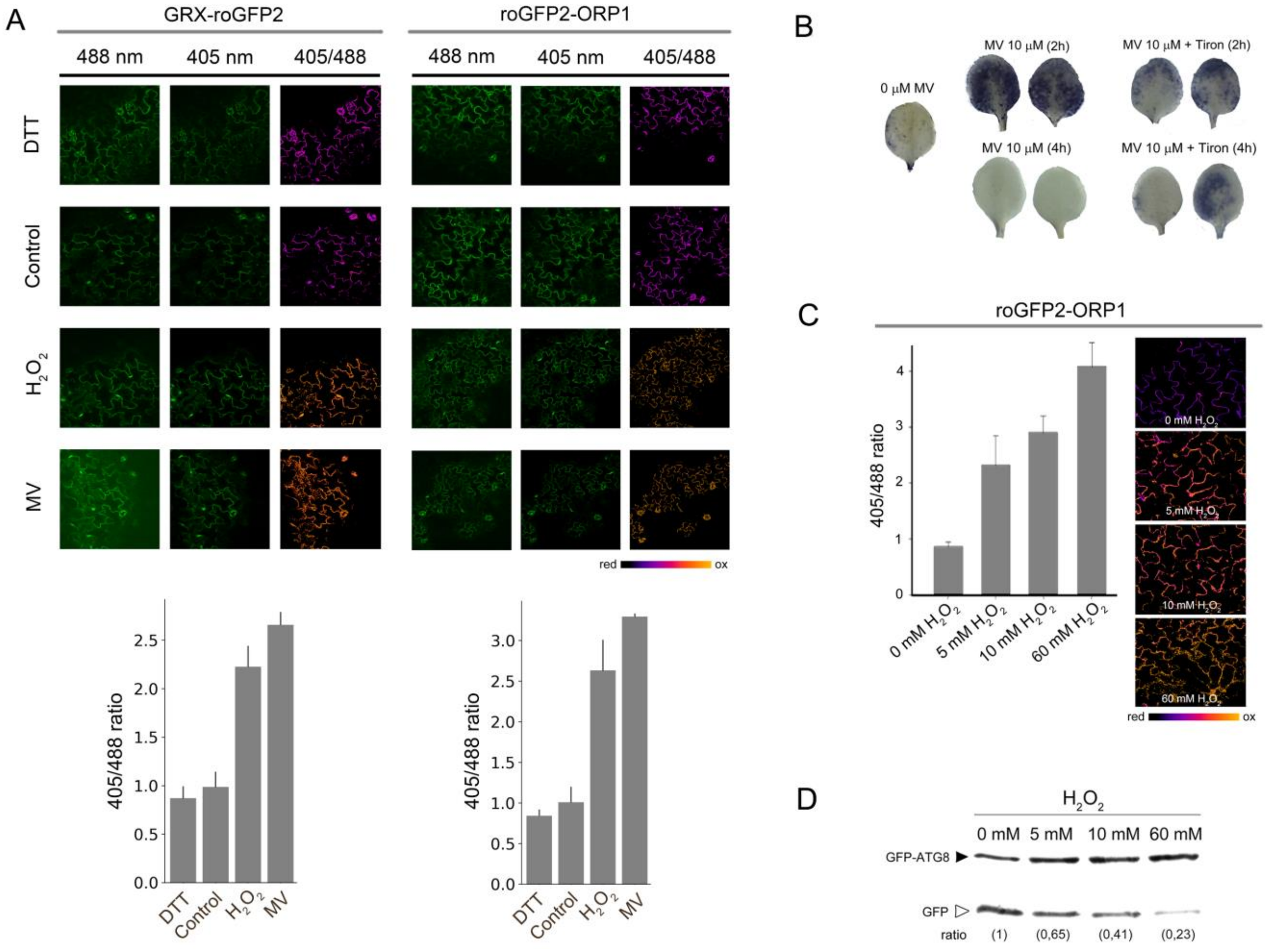


Supplementary Figure 1: Optimization of methyl viologen (MV) and $H_2O_2$ concentrations, and calibration of the redox sensors. (A) Dynamic range of the redox sensors (GRX-roGFP2 and roGFP2-ORP1) using 10 mM DTT, 10 mM $H_2O_2$ and 1µM MV. Images obtained with both lasers (488 nm and 405 nm) and the ratio between them (405/488) are observed in the different columns. For pseudocolor display, the ratio was coded on a spectral color scale ranging from blue (fully reduced) to red (fully oxidized). Representative images are shown. Graph bars show the average of cytoplasmic redox state expressed as the ratio of the fluorescence intensities measured at 405 and 488. (B) Seven-days-old seedling were exposed to 10 µM MV and stained with NBT. After 2 h, MV-treated plants exhibit a significant accumulation of ROS. The co-administration of tiron led to a reduction in ROS generation. After 4 h, plants treated with MV did not show ROS production, indicating cell death. However, co-administration of tiron conferred resistance to the oxidative treatment, as evidenced by the observed ROS production. (C) Cytoplasmic oxidation of the roGFP2-ORP1 sensor probe in Arabidopsis seedlings exposed to different $H_2O_2$ concentrations. The stress treatments were performed for 6 h, followed by the evaluation of the cytoplasmic redox state by confocal microscopy. For pseudocolor display, the ratio was coded on a spectral color scale ranging from blue (fully reduced) to red (fully oxidized). Representative images are shown. Graph bars show the average of cytoplasmic redox state expressed as the ratio of the fluorescence intensities measured at 405 and 488. (D) Autophagic flux evaluated through the degradation of the GFP-ATG8a fusion protein by western blot. Seven-day old 35S:GFP-ATG8a expressing plants were exposed to different $H_2O_2$ concentrations. Total protein (40 µg) was subjected to immunoblot analysis with anti-GFP antibodies. The GFP-ATG8a fusion has a predicted molecular weight of approximately 40 kDa (closed arrowheads); free GFP has a predicted molecular weight of 27 kDa (open arrowheads). Representative images are shown. Ratio: the numbers in brackets denote the GFP/GFP-ATG8 ratio relativized to the condition without oxidative treatment (to 0 mM $H_2O_2$).

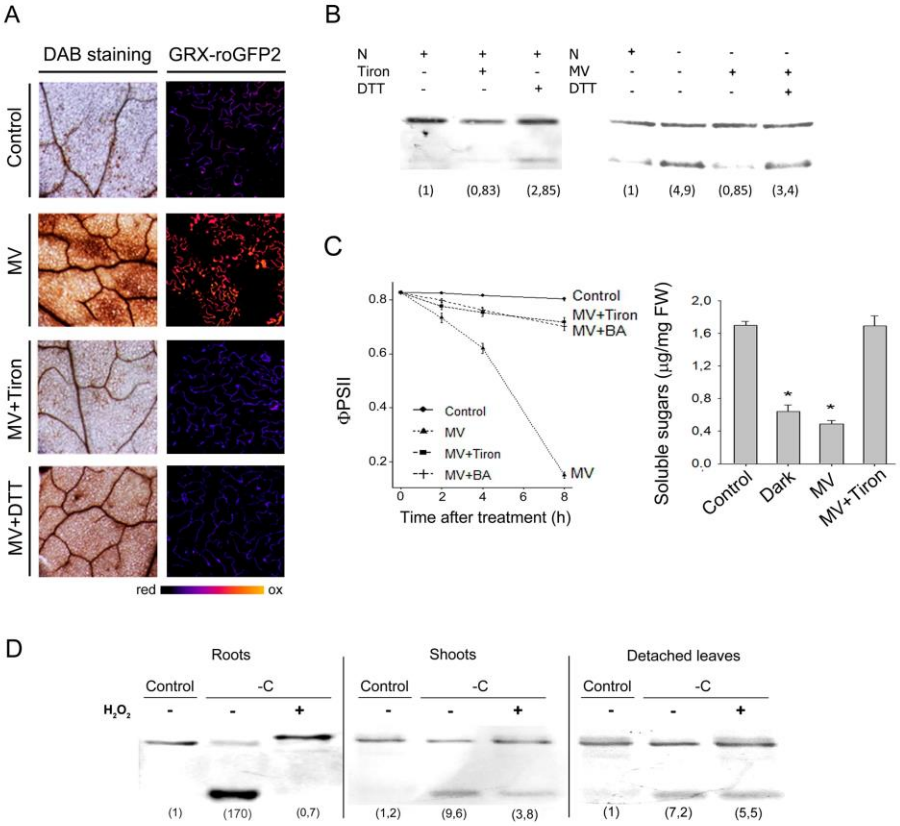


Supplementary Figure 2: Effects of DTT and tiron on (A) ROS production and cytoplasmic oxidation of the GSH:GSSG sensor probe (GRX-roGFP2), (B) autophagic flux, (C) quantum efficiency of PSII photochemistry (ΦPSII) and soluble sugar content in Arabidopsis plants exposed to 1 µM MV. (A) Production of $H_2O_2$ revealed by *in situ* staining with 3′,3′-diaininobenzidine (DAB staining) and cytoplasmic redox state (GRX-roGFP2). GRX-roGFP column is a ratiometric image. For pseudocolor display, the ratio was coded on a spectral color scale ranging from blue (fully reduced) to red (fully oxidized). Representative images are shown. (B) Autophagic flux in 7-day old seedlings evaluated through the degradation of the GFP-ATG8a fusion protein by western blot. Total protein (40 µg) was subjected to immunoblot analysis with anti-GFP antibodies. The numbers in brackets denote the GFP/GFP-ATG8 ratio relativized to the control condition. N: nitrogen; MV: methyl viologen; DTT: dithiothreitol. DTT treatments promoted autophagic flux (C) Seven-day old Arabidopsis seedlings were exposed to 1 µM methyl viologen (MV), MV + 10 mM Tiron, MV + 10 mM BA, and control condition (without MV) for 8 h, and the different physiological parameters were analyzed. (D) Inhibitory effects of 10 mM $H_2O_2$ on autophagic flux were analyzed in roots and shoots of 2-week seedlings. Plants were subjected to 16 h of darkness and treated with 10 mM $H_2O_2$, with root and shoot tissues sampled separately for analysis. Furthermore, the effects of 10 mM $H_2O_2$ were also assessed in detached leaves of 28-days old plants. Total protein (40 µg) was subjected to immunoblot analysis with anti-GFP antibodies. The numbers in brackets denote the GFP/GFP-ATG8 ratio relativized to the control condition.

**Sequence alignment of ATG4 proteases from different organisms**
Colored by: identity

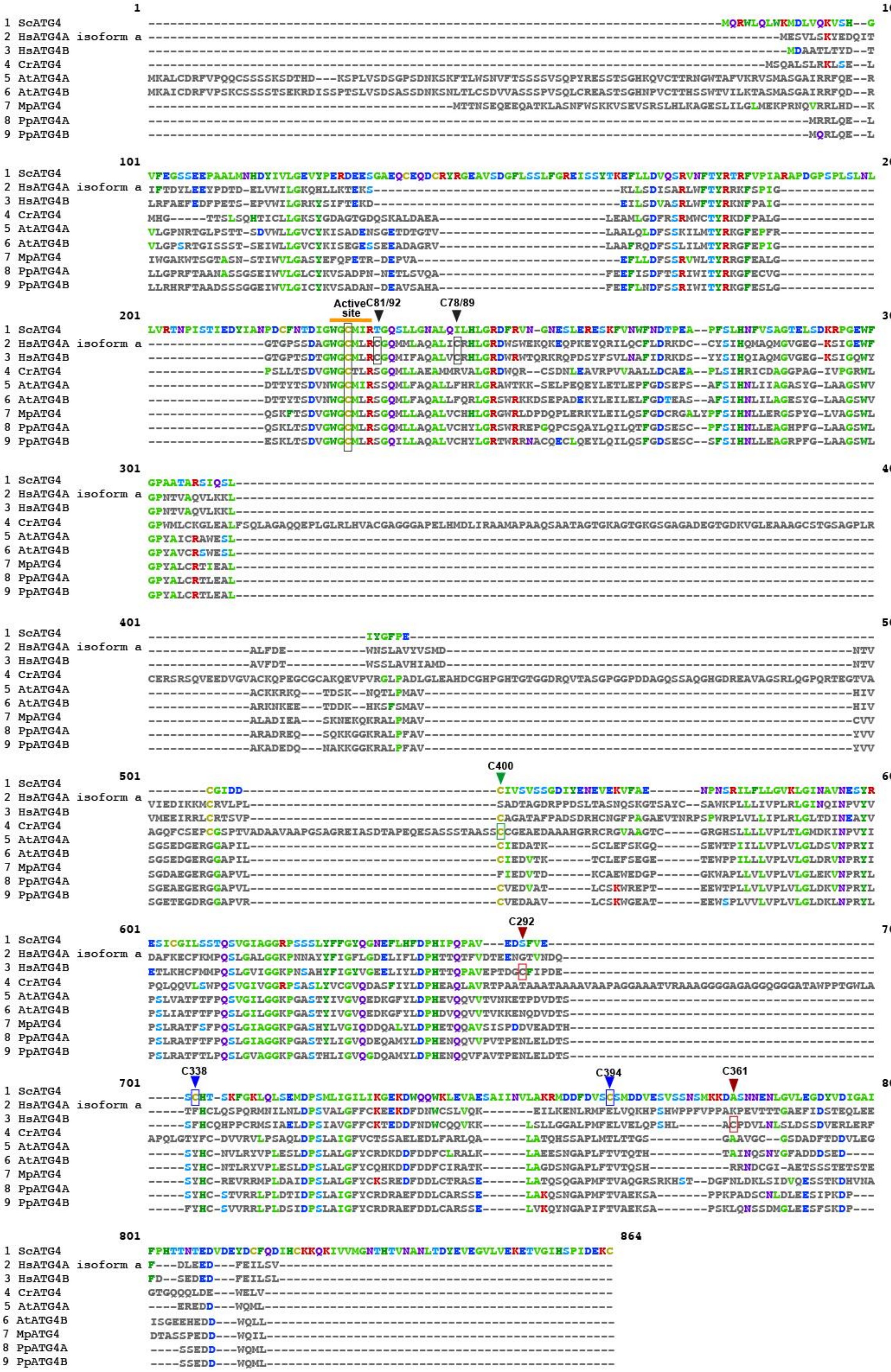

Supplementary Figure 3: Sequence alignment of ATG4 proteases from different organisms including *Saccharomyces cereviseae* (QHB11163.1), *Homo sapiens* (ATG4A- NP_001308219.1, ATG4B, NP_847896.1) *Chlamydomonas reinhardtii* (Cre12.g510100.t1.1), *Arabidopsis thaliana* (ATG4A- AT2G44140.1, ATG4B- AT3G59950.1), *Marchantia polymorpha* (Mapoly0153s0038.1.p) and *Physcomitrium patens* (ATG4A- Pp3c11_22820V3.1.p, ATG4B, Pp3c7_280V3.1.p). The conserved active site (WGCT/ML/IR) of these proteases, including the catalytic Cys77 (number according to HsATG4A) which is essential for ATG4 activity (Tanida et al., 2004) is highlighted in orange. Cysteine residues shown to be redox sensitive-sites are indicated in different colors: Cys400 in Chlamydomonas (green), Cys338 and Cys394 in yeast (blue), Cys292 and Cys361 in HsATG4B (red). Additionally, HsATG4A and HsATG4B contain two conserved cysteine residues Cys81 and Cys92 (HsATG4A), or Cys78 and Cys89 (HsATG4B), marked with black arrows. These residues are not conserved in yeasts or other eukaryotes as algae or plant ATG4s. $H_2O_2$, at the site of Cys81/Cys78, is an oxidative signal that inactivates the delipidation activity of ATG4 at the site of autophagosome formation, promoting lipidation of ATG8. Interestingly, the cysteine residue corresponding to Cys400 in Chlamydomonas is conserved in yeast, human ATG4B, and both Arabidopsis and Physcomitium ATG4 isoforms.

**A) Sequence alignment of ATG3 proteins from different organisms**

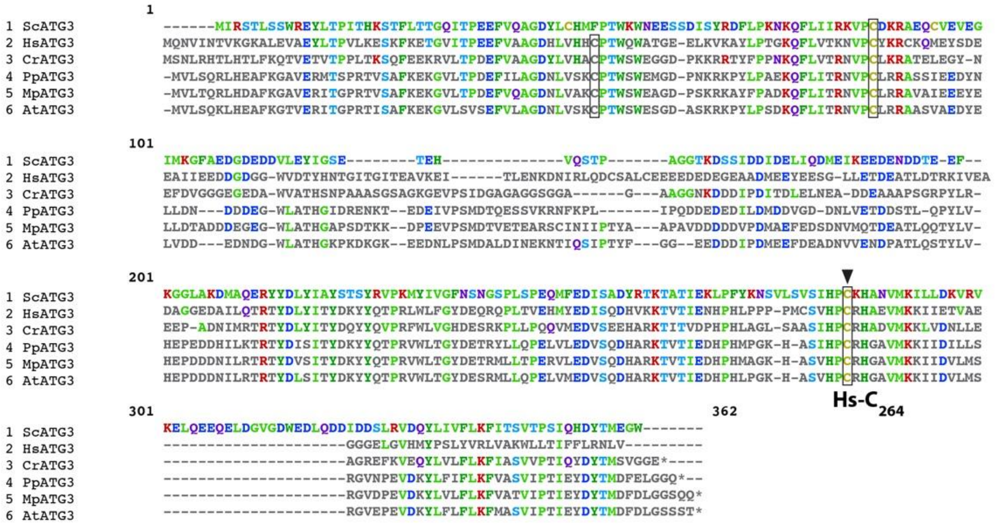


**B) Sequence alignment of ATG7 proteins from different organisms**

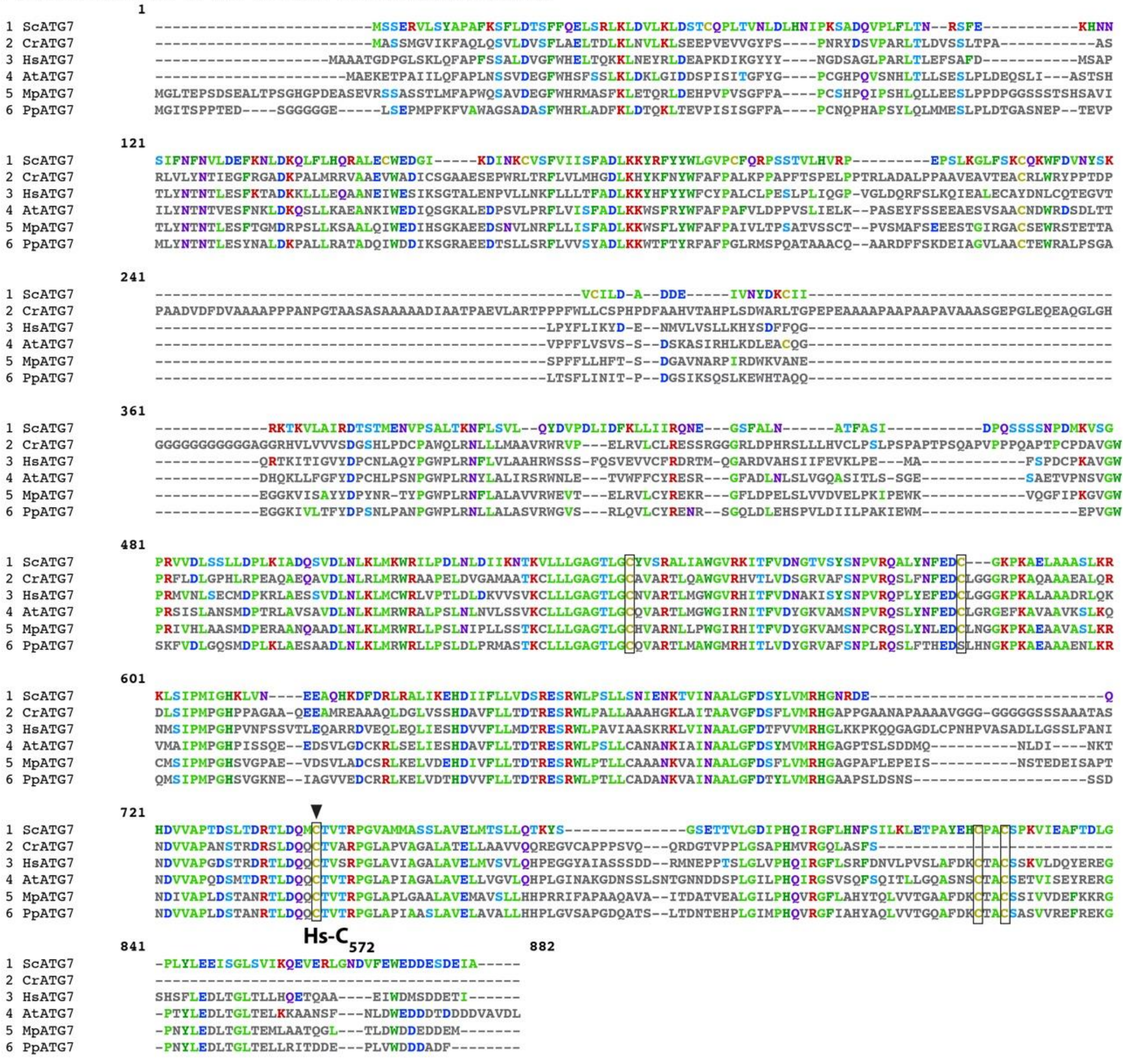

Supplementary Figure 4: (A) Sequence alignment of ATG3 proteins from different organisms including *Saccharomyces cerevisiae* (QHB11385.1), *Homo sapiens* (NP_001265641.1), *Chlamydomonas reinhardtii* (Cre02.g102350_4532.1), *Arabidopsis thaliana* (AT5G61500.1), *Marchantia polymorpha* (Mapoly0003s0208.1.p) and *Physcomitrium patens* (Pp3c8_11900V3.1.p). The catalytic Cys264 (number according to HsATG3) is conserved in all proteins studied (black box with a black arrow). (B) Sequence alignment of ATG7 proteins from different organisms including *Saccharomyces cerevisiae* (QHB09162.1), *Homo sapiens* (NP_001336161.1), *Chlamydomonas reinhardtii* (Cre03.g165215.t1.1), *Arabidopsis thaliana* (AT5G45900.1), *Marchantia polymorpha* (Mapoly0015s0071.1.p) and *Physcomitrium patens* (Pp3c24_8100V3.1.p). The catalytic Cys572 (number according to HsATG7) is conserved in all proteins studied (black box with a black arrow). Under non-reducing conditions, ATG3 and ATG7 form stable thioester complexes with LC3 (ATG8) to remain active. However, under pro-oxidizing conditions, thiol oxidation disrupts these complexes, resulting in an intermolecular disulfide bond between Cys264 (ATG3) and Cys572 (ATG7), inhibiting ATG8 lipidation (Frudd et al., 2018). Other conserved cysteines (black boxes) may also warrant further study.

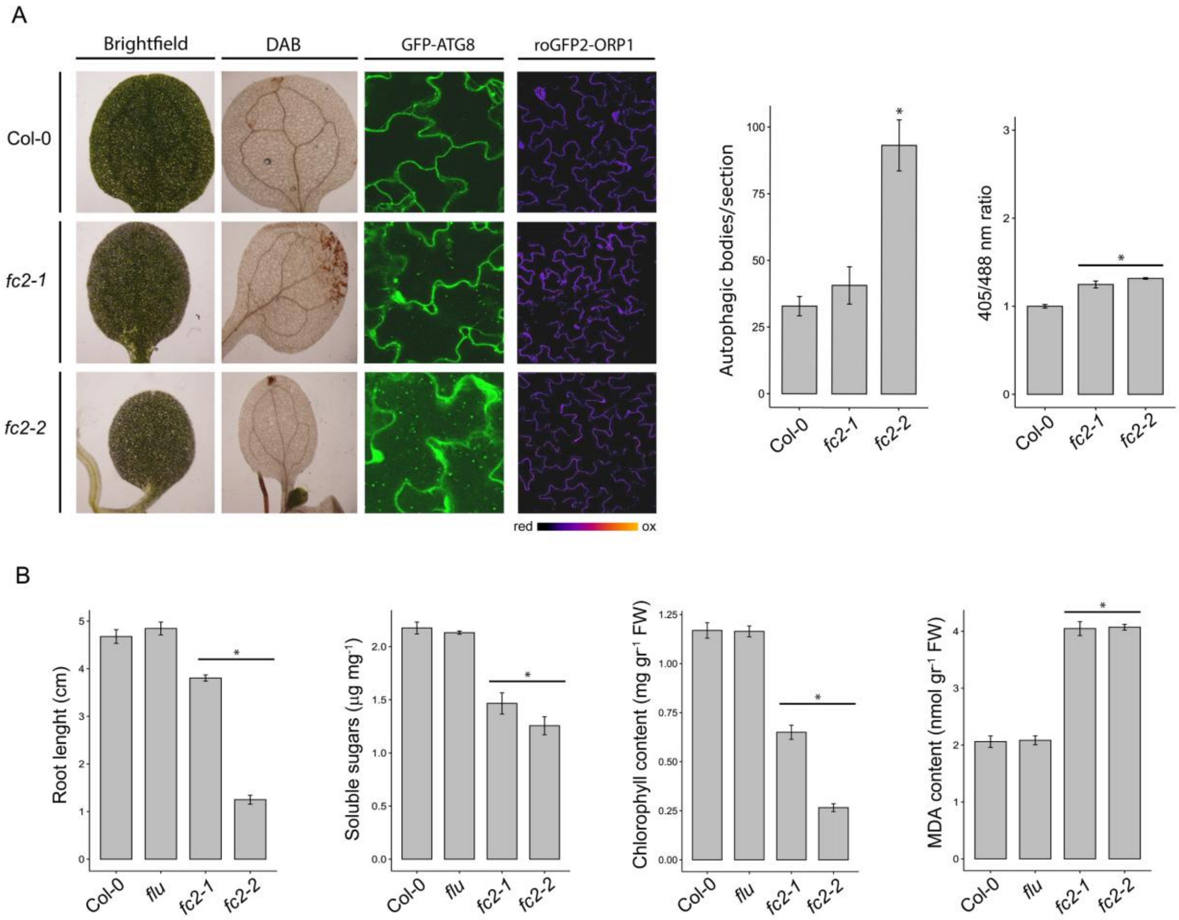


Supplementary Figure 5: Characterization of *fc2* and *flu* mutant plants under continuous light conditions (control conditions). Wild type (Col-0), *fc2* and *flu* mutant plants were grown 7 days under continuous light conditions and different growth and physiological parameters were evaluated. (A) Production of $H_2O_2$ was revealed by *in situ* staining with 3′,3′-diaininobenzidine (DAB); autophagic bodies accumulation (GFP-ATG8 columns) and the cytoplasmic redox state (GFP2-Orp1 column) analyzed by confocal microscopy. To induced autophagic bodies accumulation (GFP-ATG8 column), MS liquid medium was supplemented with 1 µM Concanamycin A. GFP2-Orp1 column is a ratiometric image. For pseudocolor display, the ratio was coded on a spectral color scale ranging from blue (fully reduced) to red (fully oxidized). Representative images are shown. At least 4 plants were analyzed. Bar plots show the average number of autophagic bodies and the average of cytoplasmic redox state. The average of autophagic bodies number was calculated from single-plane confocal images. Two sections from at least 4 plants were analyzed. The average of cytoplasmic redox state is expressed as the ratio of the fluorescence intensities measured at 488 and 405 (data expressed relative to Col-0). At least 4 plants were analyzed. (B) Average root length, soluble sugar, chlorophyll and MDA content in the different genotypes after growing seven days under continuous light conditions (*$P < 0.05$; error bars: means ± SEM, DGC test).

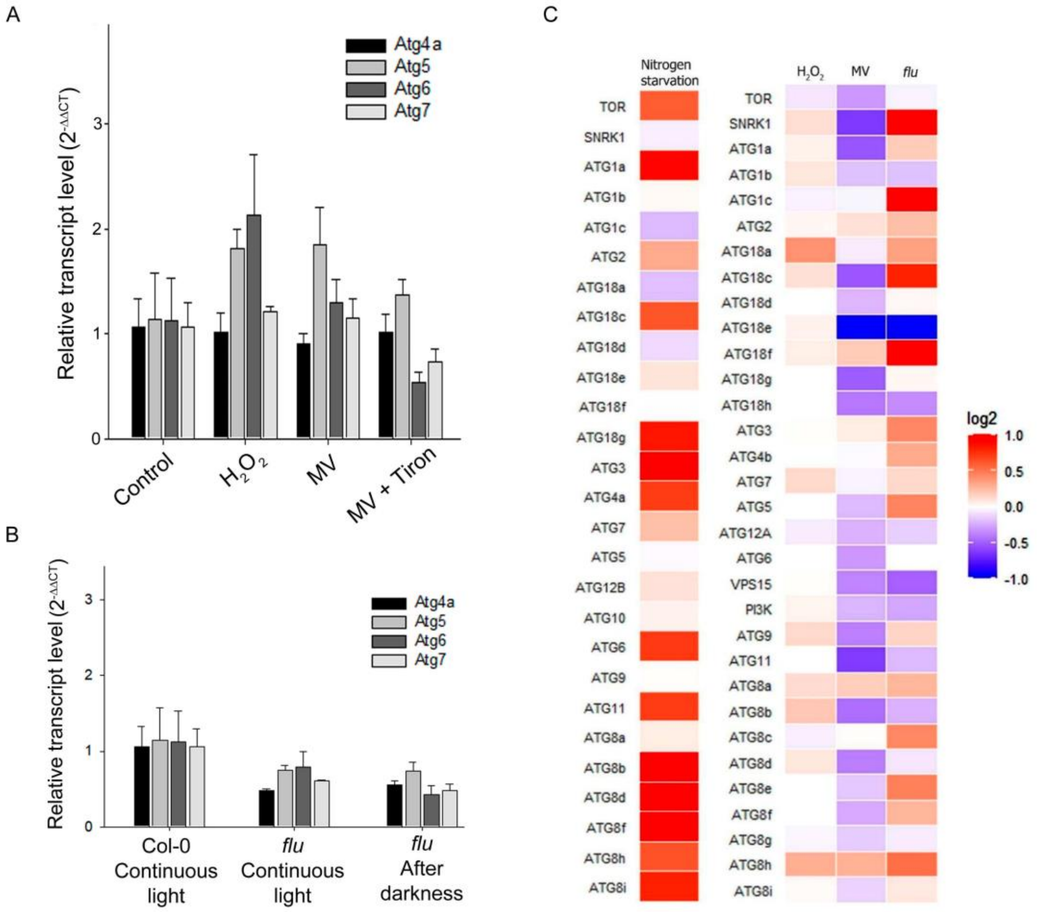


Supplementary Figure 6: Transcript level of *Atg* genes in Arabidopsis plants subjected to different oxidative treatments. (A) Seven-day old wild type Arabidopsis plants (Col-0) were exposed to oxidative stress conditions for 4 h (10 mM $H_2O_2$, 1 µM methyl viologen (MV), 1 µM methyl viologen + 10 mM tiron (MV + Tiron)), followed by the analysis of the transcript levels of *Atg* genes through RT-qPCR. Data are relative to control condition (assigned a value of 1). (B) Seven-day old *flu* mutant plants grown under continuous light conditions were exposed to darkness for 8 h and re-exposed to light for 8 h or kept in continuous light (control), and the expression of Atg genes were analyzed by RT-qPCR. Data are relative to Col-0 genotype (assigned a value of 1). (C) Publicly available microarray data were utilized to compare the expression of *Atg* genes under various conditions, including nitrogen deficiency (Rubin *et al.*, 2009), treatment with 10 mM $H_2O_2$ for 24 h (Willems *et al.*, 2016), treatment with 10 µM MV for 2 h (Scarpeci *et al.*, 2008), and the examination of *flu* mutant plants subjected to an 8-hour cycle of darkness followed by 2 h of light (Op Den Camp *et al.*, 2003). Expression values were normalized to the corresponding control conditions employed in each study and were expressed as $log_2$ values.
